# A Hybrid Transistor with Transcriptionally Controlled Computation and Plasticity

**DOI:** 10.1101/2023.08.16.553547

**Authors:** Yang Gao, Yuchen Zhou, Xudong Ji, Austin J. Graham, Christopher M. Dundas, Ismar E. Miniel Mahfoud, Bailey M. Tibbett, Benjamin Tan, Gina Partipilo, Ananth Dodabalapur, Jonathan Rivnay, Benjamin K. Keitz

## Abstract

Organic electrochemical transistors (OECTs) are ideal devices for translating biological signals into electrical readouts and have applications in bioelectronics, biosensing, and neuromorphic computing. Despite their potential, developing programmable and modular methods for living systems to interface with OECTs has proven challenging. Here we describe hybrid OECTs containing the model electroactive bacterium *Shewanella oneidensis* that enable the transduction of biological computations to electrical responses. Specifically, we fabricated planar p-type OECTs and demonstrated that channel de-doping is driven by extracellular electron transfer (EET) from *S. oneidensis*. Leveraging this mechanistic understanding and our ability to control EET flux via transcriptional regulation, we used plasmid-based Boolean logic gates to translate biological computation into current changes within the OECT. Finally, we demonstrated EET-driven changes to OECT synaptic plasticity. This work enables fundamental EET studies and OECT-based biosensing and biocomputing systems with genetically controllable and modular design elements.

## Introduction

Devices that transduce and amplify biological and chemical activity into electrical signals are highly desirable in a number of fields including sensing^1,2^, neuromorphic computing^3^, cellular computing^4^, and wearable electronics^5^. For several of these applications, organic electrochemical transistors (OECTs) have emerged as ideal devices owing to their use of aqueous electrolytes, compatibility with biological systems, and low operating voltages^6,7^. In contrast to conventional electronics that rely on semiconducting and dielectric materials, OECTs utilize ions within an electrolyte to alter the doping state and conductivity of an organic mixed ionic-electronic conducting channel^8^. Because the entire volume of the channel is accessible to ions in the electrolyte, a relatively small potential change at the gate can significantly alter the channel’s conductivity, giving OECTs exceptional transconductance and sensitivity. In addition to sensing and flexible electronics applications, OECTs are promising devices for neuromorphic computing because synaptic weight, usually defined as channel conductance, can be altered by controlling ion transport in and out of the channel^9,10^. Overall, the inherent ability of OECTs to couple ionic and electronic transport makes them ideal devices for merging aspects of biological and traditional computation.

While ionic diffusion into the channel is typically controlled using an applied voltage at the gate electrode, biological or reduction-oxidation (redox) reactions in the electrolyte can also change the channel doping state. In effect, redox reactions can function as a secondary gate in the OECT. For example, lipid bilayer functionalization of the channel or gate followed by insertion of gated ion channels, nanobodies, or other biomolecules allow OECTs to sense a variety of chemical and biological stimuli^11,12^. Similarly, redox-active enzymes, such as lactate or glucose oxidase, can directly transfer electrons into the channel to tune its conductivity in response to chemical substrates^13^. These applications highlight the usefulness of OECTs for sensing and diagnostic applications, but achieving more complex sensing is challenging because individual enzymes, proteins, and other biomolecules are only capable of limited computation on a single device. In contrast, living cells perform a variety of extremely complex and robust computations that could potentially be tied to an OECT output. For example, bacteria can be engineered to perform computations including Boolean logic operations^14,15^, analog/digital signal processing^16^,cellular computing^4^, and neuromorphic computing^17^. While OECTs have long been used to detect bacteria or sense the presence of specific metabolites^18^, coupling more advanced computations to an electrical output via genetic circuits that regulate protein expression, small molecule synthesis, or other outputs that reliably interface with an OECT has proven challenging.

One promising strategy for interfacing bacterial computation with OECTs and other electronic devices is the use of electroactive bacteria. While all bacteria regulate ion flux into and out of the cell, electroactive bacteria can directly modulate electron transport across the cellular membrane in a process known as extracellular electron transfer (EET). Under anaerobic conditions, electroactive bacteria couple their central carbon metabolism to the oxidation or reduction of metal species in the environment via EET. Although naturally-occurring metals and metal oxides are the most well-studied electron acceptors for EET, synthetic materials including nanoparticles^19^, conducting polymers^20^, and a variety of electrode materials can also accept electron flux from these bacteria^21^. The use of electroactive bacteria in microbial fuel cells and similar bioelectronic devices has been extensively studied for power generation applications^22^. More recently, advances in synthetic biology and the engineering of model electroactive bacteria, such as *Shewanella oneidensis* and *Geobacter sulfurreducens,* have enabled broader applications of EET in sensing and biological computation^23,24^. To facilitate this transition, we, and others, have created genetic circuits that tightly regulate the expression of EET-relevant genes, allowing EET flux to be turned on and off in response to specific combinations of chemical, biological, and physical stimuli^25,26^. Thus, genetic regulation over electron transport via EET has the potential to serve as a universal interface between bacteria and electronic devices, including OECTs. Electrochemical transistors inoculated with electroactive bacteria can be conceptualized as dual-gate devices, wherein the initial gate represents the conventional electrode. The modulation induced by the second gate arises from electrochemical interactions facilitated by the bacterial cells. These interactions influence the charge balance within the channel, consequently altering its doping states. Despite this promise, instances of directly coupling electroactive bacteria with OECTs remain limited. Méhes et al. presented a notable example of real-time cellular EET activity monitoring with p-type OECT^18^. Using *S. oneidensis* (wild-type strain, MR-1) and the inherent amplification of OECT, changes in cellular metabolism beyond the limit of the conventional electrochemical setups were detected. While this work established an important proof of principle that EET could be detected using OECTs, we hypothesized that developing an improved mechanistic understanding of interactions between bacteria and the OECT combined with genetic regulation over EET flux could enhance our understanding of bacteria-OECT interactions, couple biological computation to an electrical response, and drive programmable changes to OECT synaptic plasticity.

Here, we developed hybrid transistors consisting of genetically engineered electroactive bacteria in planar p-type poly(3,4-ethylenedioxythiophene):poly(styrenesulfonate) (PEDOT:PSS) OECTs. Using *S. oneidensis* as a model EET-capable species, we first determined that channel conductance can be altered via the number and metabolic state of cells growing in the electrolyte. Next, we unraveled the biological and chemical mechanisms responsible for de-doping and current changes within the PEDOT:PSS channel using a combination of electrochemical and spectroscopic methods. To further illuminate the de-doping mechanism, we deployed genetically engineered *S. oneidensis* strains to regulate EET flux and analyzed the corresponding OECT outputs. Leveraging this mechanistic information, we converted EET flux from mutant strains carrying genetic Boolean logic circuits to electrical readouts, allowing the OECT to detect complex combinations of environmental signals. Finally, we characterized the synaptic behaviors of hybrid OECTs containing *S. oneidensis* strains, showcasing tunable synaptic weight tied to transcriptional outputs. Overall, our work establishes a new foundation for biosensing and biocomputing by augmenting OECT performance with genetically controllable inputs.

## Results

### Device Fabrication and Initial Characterization

In contrast to most bioelectrochemical systems, such as microbial fuel cells^27^, OECTs require microfabrication and rigorous spatial control over conducting polymer deposition. PEDOT:PSS is frequently used for OECT channels due to its high biocompatibility, chemical/physical stability, and high electrical conductivity^6^. Given these advantages and previous studies examining *S. oneidensis* in the presence of conductive polymer electrodes^20^, we selected PEDOT:PSS as the OECT channel material. We fabricated planar OECTs on quartz microscope slides with Ti/Au as the gate, source, and drain electrodes (Figure 1a and 1b, Figure S1). The channel region and the tip of the gate surface were coated with PEDOT:PSS. Following fabrication, we first verified that *S. oneidensis* MR-1 could grow in the electrolyte (Shewanella Basal Medium, Table S2) and colonize the OECT under anaerobic conditions. OECT electrodes were constantly biased at V_GS_ = 0.2 V and V_DS_ = −0.05 and fluorescence microscopy images were taken 24 hours post-inoculation to determine cell viability and distribution within the device (Figure 1c and 1d, Extended Data Figure 1a and 1b). Cells maintained a viability of 66.8 ± 13.8 % and were found near the gate, channel, source, drain, and spaces in between (Extended Data Figure 1c and 1d). To corroborate these results, we monitored colony forming units (CFUs) and optical density at 600 nm (OD_600_) within the OECT electrolyte. CFU counts were measured 24 hours post-inoculation in OECTs under constant V_DS_ = −0.05 V and V_GS_ at −0.5 V, −0.2 V, 0.0 V, and 0.2 V. The OD_600_ readings were measured in OECTs under constant V_DS_ = −0.05 V and V_GS_ = 0.2 V. Consistent with our cell viability measurements, CFU counts dropped slightly after 24 hours and showed no dependence on the gate potential (Figure 1e). Similarly, OD_600_ readings remained consistent during OECT operation, indicative of minimal cell growth over a 24-hour period (Extended Data Figure 1f). When fumarate was included as a soluble electron acceptor to support anaerobic growth, cell viability improved to 80.2 ± 10.4% after 24 hours post-inoculation (Extended Data Figure 1c and 1e). As expected, the presence of fumarate also resulted in significant cell growth within the OECT, as indicated by increased CFU counts and OD_600_ values (Extended Figure 1g and 1h). Taken together, these results indicate that PEDOT:PSS in the OECT can support cell maintenance, but not robust growth. While fumarate facilitated cell growth, we did not include it in subsequent experiments as its presence could discourage *S. oneidensis* from interacting with the PEDOT:PSS.

**Figure 1.**
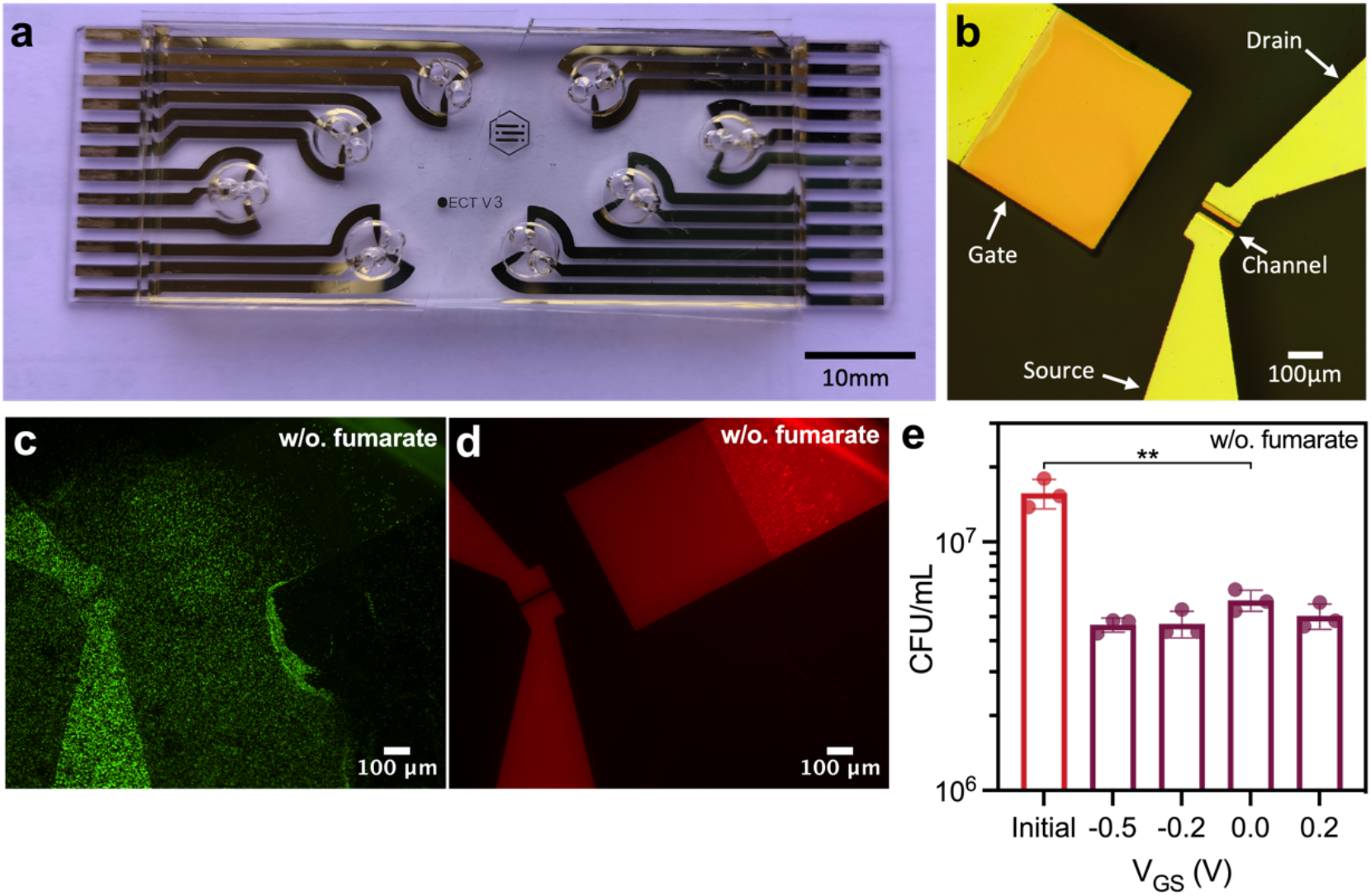
Photo and microscopy images of the OECT. (**a**) 8 OECTs on a microscope slide with PDMS sheets to form the OECT chambers, and (**b**) top view of a single OECT. (**c, d**) Fluorescence microscopy images of cells stained with LIVE/DEAD^®^ BacLight™ cell assay with (**c**) live cells shown in green, and (**d**) dead cells shown in red. Background fluorescence signals from the gold electrodes (red channel) are also visible. Cells were supplemented with 20 mM lactate and no additional electron acceptor. (**e**) Colony forming units (CFUs) were counted 24 hours after OECTs operation with constant V_DS_ = −0.05 V and V_GS_ biased at −0.5 V, −0.2 V, 0.0 V, or 0.2 V. Cells were supplemented with 20 mM lactate and no additional electron acceptor. Representation of p-values (n.s. p > 0.05, * p ≤ 0.05, ** p ≤ 0.01, *** p ≤ 0.001, **** p ≤ 0.0001) and data show the mean ± SD of 3 biological replicates.

Under anaerobic conditions, *S. oneidensis* transfers electrons outside of the cell through the metal-reducing (Mtr) pathway, which is composed of three proteins MtrC, MtrA, and MtrB encoded on a single operon^28^ (Figure 2a). Based on previous work examining *S. oneidensis* on PEDOT:PSS coated electrodes^20^, we predicted that the bacteria could potentially influence channel current in the OECT through two major mechanisms (Figure 2b). First, at positive gate voltages that energetically favor extracellular electron transfer, electrons may flow from bacteria cells to the gate. To maintain electrical neutrality, electrons will travel to the source via the external circuit. Subsequently, the electrons will flow out of or accumulate on the source electrode, where they can combine with PEDOT^+^ and de-dope the PEDOT:PSS channel. Alternatively, if the reduction potential of the channel is higher than that of the cell, the bacteria can directly reduce and de-dope the PEDOT:PSS, even in the absence of an applied potential at the gate. In either mechanism, biologically-driven de-doping of the PEDOT:PSS channel should be characterized by a pronounced decrease in channel current over time. Indeed, we found that devices inoculated with *S. oneidensis* MR-1 showed decreased channel current within 30 minutes relative to abiotic controls (Figure 2c). Next, to examine device stability, OECTs were exposed to oxygen after 48 hours of operation in the presence of *S. oneidensis*. The I_DS_ recovered close to the original levels (Extended Data Figure 2a and 2b). To further examine material stability within the device, OECTs containing *S. oneidensis* were gently washed with soapy water and examined with atomic force microscopy (AFM). We found no significant changes in PEDOT:PSS film thickness compared to pre-inoculation films (Figure S2a). Similarly, the channel surface roughness exhibited minimal alteration, with root mean square (RMS) roughness values of 2.4 nm and 2.6 nm for pre-inoculation and post-inoculation films, respectively (Figure S2c and S2d). The PEDOT grain sizes were extracted from phase images (Figure S2e and S2f), and subtle segregation of the PEDOT cores (brighter color) was discernible from the post-inoculation samples with PSS filling the space in between (darker color)^29^. Importantly, the histograms for PEDOT grain size exhibited comparable distributions, suggesting minimal changes in the material constitution following bacterial incubation (Figure S2b). Collectively, these results indicate good device stability and a reversible mechanism between *S. oneidensis* and OECT channel de-doping. Finally, to facilitate comparisons between different biological conditions and account for dynamic changes to channel current, the rate of current decay was fitted to a single exponential decay model to extract a rate constant characteristic of EET-driven channel current decreases (Figure 2c).

**Figure 2.**
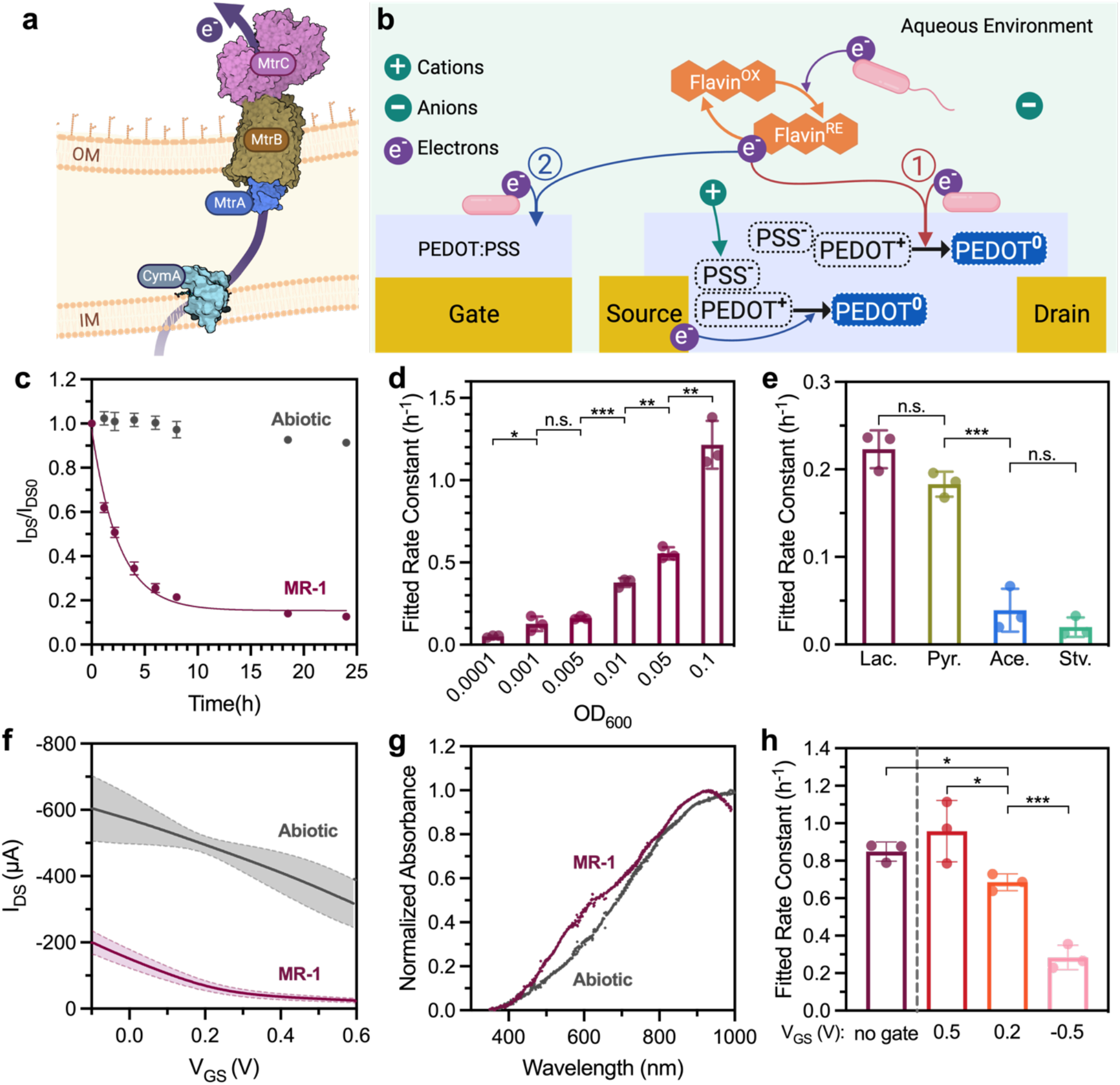
The proposed cellular de-doping mechanisms and OECT output changes induced by *S. oneidensis* MR-1. (**a**) Illustration of the MtrCAB pathway directing electrons to extracellular electron acceptors. (**b**) Proposed channel de-doping mechanisms through direct and mediated EET. **Path 1**, cellular metabolic electrons are transferred directly to the PEDOT:PSS channel. **Path 2**, electrons are transferred to the channel via the gate and external circuits (circuits not shown). (**c**) The I_DS_/I_DS0_ curve for OECTs inoculated with *S. oneidensis* MR-1, cells were supplemented with 20 mM sodium lactate and no additional electron acceptors. Initial inoculum OD_600_ = 0.01. (**d, e**) Fitted rate constants for *S. oneidensis* inoculated OECT with different (**d**) inoculation densities (OD_600_) or (**e**) carbon sources. Samples with different inoculation densities were supplemented with 20 mM sodium lactate as the carbon source. Carbon source and starvation samples were prepared at an inoculation OD_600_ of 0.01 and supplemented with either 20 mM sodium lactate (Lac.), 20 mM sodium pyruvate (Pyr.), or 20 mM sodium acetate (Ace.). No carbon source was added to the starved cells (Stv.). (**f**) Transfer curves were measured in the presence and absence of *S. oneidensis*. The shaded region indicates the range of standard deviation. (**g**) UV-Vis spectra of the PEDOT:PSS channel in the presence and absence of *S. oneidensis*. (**h**) I_DS_ decay profiles compared for OECTs biased at different gate voltages and gate-removed 2-electrode devices. Representation of p-values (n.s. p > 0.05, * p ≤ 0.05, ** p ≤ 0.01, *** p ≤ 0.001, **** p ≤ 0.0001) and data show the mean ± SD of 3 biological replicates.

On a per-cell basis, currents from EET flux are relatively small^30^. However, more cells should generate more flux and faster response times. As expected, the measured rate constant associated with the decrease in OECT current was proportional to the size of the starting cell population (Figure 2d, Extended Data Figure 2c). Because EET flux is connected to the bacteria cells’ central carbon metabolism, different carbon sources generate varying amounts of EET flux. Lactate is the preferred carbon source for *S. oneidensis*, followed by pyruvate, while acetate cannot be metabolized under anaerobic conditions^31^. As predicted, cells metabolizing lactate or pyruvate yielded the fastest current response while starved cells or those metabolizing acetate exhibited rates closer to abiotic controls (Figure 2e, Extended Data Figure 2d). These results are strong indicators that cells remain metabolically active within the OECT and that cellular metabolic flux is correlated with channel de-doping. Furthermore, to confirm the channel current reduction is driven by live *S. oneidensis* MR-1 cells, OECTs were inoculated with heat-killed and lysed *S. oneidensis* MR-1 cells, cell metabolic products from the supernatant of overnight *S. oneidensis* cultures, and live *E. coli* MG1655 cells that are incapable of EET. As depicted in Extended Data Figure 2e and 2f, only metabolically-active and lysed *S. oneidensis* MR-1 caused significant I_DS_ decay. Relative to living cell controls, the lysed samples exhibited a marked linear and slower I_DS_ decay profile that is most likely a result of the release of redox-active components, such as thiols, NADH (nicotinamide adenine dinucleotide), quinones, cytochromes, and other reductants following cell lysis. Overall, these results demonstrate that de-doping of PEDOT:PSS and associated decreases in channel current are closely tied to cellular metabolism.

### Mechanism of OECT Channel De-doping

Depletion mode OECTs are characterized by a decrease in current as the bias voltage at the gate becomes increasingly positive. Accordingly, we measured the transfer curves of devices inoculated with bacteria and observed a noticeable decrease in the channel current and source electrode potential after 24 hours of cell incubation in the device (Figure 2f, Extended Data Figure 3a). While the applied gate and drain voltages were noted as V_GS_ and V_DS_ with respect to the source electrode, the effective gate voltages (V_G_^eff^) were determined by comparing the measured source voltages (V_S_) against an Ag/AgCl pellet pseudo-reference electrode (RE), given by V_G_^eff^ = -V_S_. The measured decrease of the source electrode potential is indicative of reductions to the source and channel, which is in accordance with our hypothesis that *S. oneidensis* MR-1 can de-dope the channel. To compare the channel current decrease and electrode potential drop, we also measured the source and gate potential against an Ag/AgCl RE with fixed V_GS_ = 0.2 V and V_DS_ = −0.05 V (Extended Data Figure 3b). The strong correlation between decreasing V_G_ and |I_DS_| (absolute) values also explains the changing I_DS_ rate as cells retain metabolic activity and continue to perform EET within the OECT.

To further verify that the observed channel current reduction in the presence of *S. oneidensis* MR-1 was due to de-doping of the PEDOT:PSS channel, we conducted *in-situ* UV-Vis measurement using a modified OECT with a larger channel area (Extended Data Figure 3c). Previous studies have shown that the doping state of PEDOT:PSS can be characterized via UV-Vis spectroscopy^32^. Specifically, the neutral and polaronic states correspond to absorption peaks around 650 nm and from 800 nm to far infrared, respectively^33,34^. We initially measured spectra in abiotic devices using a non-polarizable Ag/AgCl pellet gate biased at varying voltages. As depicted in Extended Data Figure 3d, the neutral PEDOT:PSS peak at 650 nm increased as the gate voltage became more positive and the channel was increasingly de-doped. Correspondingly, the absorbance around 900 nm for the PEDOT:PSS polaron decreased sharply as the gate bias voltage increased from 0.6 V to 1.0 V. Next, large channel OECTs containing an Au gate were inoculated with *S. oneidensis* MR-1 (inoculation OD_600_ = 0.01) and operated continuously with V_GS_ = 0.2 V and V_DS_ = −0.05 V. To account for absorption from cells, a channel-less OECT with the same inoculum and operation conditions was used as the blank before each spectrum measurement. As shown in Figure 2g and Extended Data Figure 3e, the channel UV-Vis spectrum during the low I_DS_ plateau (ca. 20.5 hours after inoculation) displayed a similar profile to the de-doped abiotic channel with an Ag/AgCl gate biased at 0.5 V, indicating the presence of *S. oneidensis* cells had a comparable de-doping effect as an Ag/AgCl gate poised at this potential.

Because these EET de-doping experiments were conducted with cells present on both the gate and channel, the direct biological reduction of the channel could not be isolated from the observed decreases in current in devices containing *S. oneidensis* MR-1. Therefore, we also examined a two-electrode version of the small OECT where the Au gate electrode was completely removed and EET can only affect the channel doping state via direct reduction (Extended Data Figure 3f). Similar to our previous continuous OECT operation conditions, a constant V_DS_ = −0.05 V was applied to the 2-electrode devices. As shown in Figure 2h and Extended Data Figure 3g, the 2-electrode devices exhibited pronounced I_DS_ decay upon inoculation with *S. oneidensis* and a decay rate comparable to our three terminal OECTs with V_GS_ = 0.2 V. The pronounced I_DS_ change in 2-electrode devices confirmed the direct and potent interaction between bacteria cells and the PEDOT:PSS channel.

De-doping of PEDOT:PSS could also occur through interaction of *S. oneidensis* with the gate and subsequent electron accumulation on the source. To further examine this possibility, we compared the channel current decay rates from our original three-terminal OECTs with different gate bias voltages. Gate voltages of 0.5 V, 0.2 V, and −0.5 V were selected, with decreasing energetic favorability for EET to the gate. As shown in Figure 2h, the I_DS_ decay rate constants increased with more positive gate bias voltages for biotic OECTs, while the abiotic OECTs showed negligible current change (Extended Data Figure 3g). The ability of gate bias potential to modulate the I_DS_ rate constant confirmed our hypothesis that electrons transferred to the gate could participate in channel reduction via the external circuit. Together, these results suggest that *S. oneidensis* MR-1 can directly de-dope the PEDOT:PSS channel via electron transfer, and the gate bias voltage plays a significant role in modulating the de-doping process. Although we were unable to completely distinguish between electron transfer at the gate and channel, the ability to tune the biological response based on the applied gate voltage highlights how multiple parameters, both biological and device-based, can control OECT performance.

### EET Drives Changes in OECT Current Output

Encouraged by our results showing *S. oneidensis* MR-1 could effectively de-dope the PEDOT:PSS channel, we investigated whether these changes could be attributed to EET. In *S. oneidensis*, EET occurs through direct or indirect means. Direct EET is mediated by cell attachment and interaction with a suitable redox-active surface by the Mtr pathway, while indirect EET is controlled by the biosynthesis and secretion of flavins, which act as soluble redox shuttles^35^. We employed several genomic deletion strains and protein expression tools to control EET flux and determine which of the above mechanisms might be contributing to the OECT response (Figure 3a). Specifically, the △*bfe* strain has decreased extracellular flavin secretion due to the deletion of bacterial flavin exporter gene *bfe*, which is critical for indirect EET^36^. The △*lysis* strain has impaired biofilm formation due to the deletion of lysis operons *SO2966* to *SO2974*^37^. Finally, both △*mtrC* and △Mtr strains lack key EET proteins and exhibit decreased EET rates^38^. While the △*mtrC* strain has the key outer membrane EET proteins MtrC, OmcA, and MtrF deleted, the △Mtr strain has additional genomic deletions of periplasmic electron carriers, namely *mtrA*, *mtrD*, *dmsE*, *SO4360*, and *cctA*^38^.

**Figure 3.**
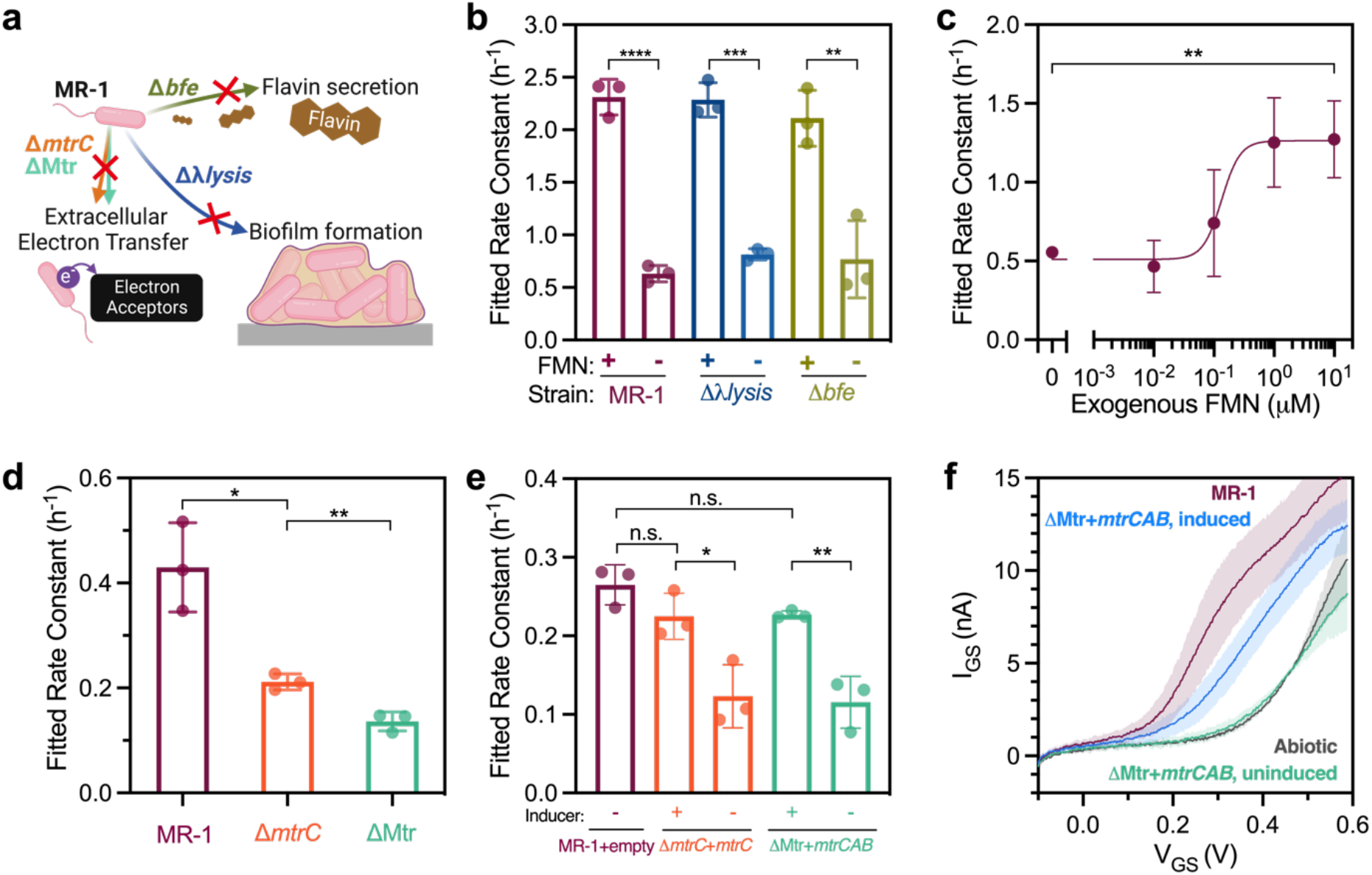
OECT response to different EET mechanisms. (**a**) Illustrations of the knockout strains with reduced secretion of FMN (△*bfe*) or impaired biofilm formation (△*λlysis*). (**b**) I_DS_ decay rate constants of knockout strains with and without 1 μM exogenous FMN. (**c**) I_DS_ decay rate constants of *S. oneidensis* MR-1 supplemented with varying concentrations of exogenous FMN. Inocula in (b) and (c) were adjusted to OD_600_ of 0.05. (**d, e**) I_DS_ decay rate constants for △*mtrC* and △Mtr knockout strains (**d**) without vector plasmids and (**e**) complemented with *mtrC* and *mtrCAB* Buffer gates. (**f**) Gate current I_GS_ from △Mtr knockout strains with induced and uninduced *mtrCAB* Buffer gates, measured at 24 hours after inoculation. The shaded region indicates the range of standard deviation. Representation of p-values (n.s. p > 0.05, * p ≤ 0.05, ** p ≤ 0.01, *** p ≤ 0.001, **** p ≤ 0.0001) and data show the mean ± SD of 3 biological replicates.

To determine the effects of the two EET pathways on OECT response, we first measured rate constants for I_DS_ decay from devices inoculated with △*bfe,* △*lysis*, and wild-type *S. oneidensis* strains (inoculation OD_600_=0.05, Figure 3b). We observed no statistically significant differences between these three strains, suggesting biofilm formation and flavin secretion are not required for PEDOT:PSS de-doping. It is likely that on the time scale of our experiments (ca. 24 hours), flavin secretion and extracellular concentration are not significant enough to affect device performance. However, adding exogenous flavins (flavin mononucleotide, 1 μM) did increase the rate of current decay for all strains tested (Figure 3b), indicating under certain conditions indirect EET can be an important contributor to OECT performance. We also observed a dose-dependent response to increasing flavin concentration. Specifically, we measured a sigmoidal response of the channel current decay rate constant with increasing exogenous FMN concentration (Figure 3c), consistent with a direct EET transfer mechanism via FMN-bound MtrC^39^. The sigmoidal relationship between flavin concentration and current decay has previously been observed in some microbial fuel cells and is likely due to flavin saturation, which is exacerbated by the small volume and low surface area of the OECT electrodes in our devices^40^.

Next, we measured current responses from the EET-deficient △*mtrC* and △Mtr strains. As expected, the rate constant and current decay rate of these strains were significantly lower than those of *S. oneidensis* MR-1, indicating that impaired EET pathways significantly slowed the channel de-doping process (Figure 3d, Figure S3c). To further confirm that differences in channel de-doping were due to EET, we constructed plasmid vectors encoding key EET genes to complement EET-deficient mutants. Specifically, we constructed an *mtrC* Buffer gate by placing the *mtrC* gene downstream of the P_tacsymO_ promoter and transformed it into the △*mtrC* strain (Figure S3d, insert). For this circuit, the presence of IPTG [IPTG = Isopropyl β-D-1-thiogalactopyranoside] allows RNA polymerase to transcribe the *mtrC* gene and ultimately increase EET flux. We also constructed a *mtrCAB* Buffer gate regulated by OC6 [OC6 = 3-oxohexanoyl-homoserine lactone], which was transformed into the △Mtr mutant (noted as △Mtr*+mtrCAB*). Mutants harboring empty vector controls (noted as +empty) without the respective EET genes were used as negative controls. Prior to inoculation into OECTs, all strains were anaerobically induced for 18–24 hours with 1mM IPTG or 100 nM OC6 to ensure steady-state protein expression. As illustrated in Figure 3e, mutants with restored EET gene expression yielded significantly higher current decay rate constants relative to empty vector controls. The lower rate constants observed in the induced samples compared to wild-type *S. oneidensis* MR-1 are likely due to the absence of other outer-membrane EET proteins, such as OmcA and MtrF, in these genetic constructs. To further verify the electrochemical activity of the induced Buffer gates samples, gate currents were measured with gate voltage scans. As shown in Figure 3f, △Mtr strains with induced *mtrCAB* exhibited higher oxidation currents compared to the uninduced control, which was almost identical to the abiotic baseline. Together, these data indicate that de-doping of PEDOT:PSS and the resulting current decreases are attributable to EET and specific EET-relevant protein expression.

### Interfacing Genetic Logic Gates with OECTs

Transistors, including OECTs, require circuit connections to perform more complex logic. In contrast, bacteria have been programmed to exhibit logic-based responses^41^, analog computing^4,16^, memory, and neuromorphic behavior^17^. For these applications, the desired computation is genetically encoded within each bacterium and the calculation is performed by the community. To determine whether similar genetic circuits could be connected to OECT performance and enable more complex logic on a single device, we evaluated transcriptionally-controlled Boolean logic gates that regulate EET gene expression and associated flux in response to combinations of small molecule inputs. Specifically, we leveraged previously developed NAND and NOR transcriptional logic gates that control EET flux in response to combinations of inducer stimuli^25^. As building blocks in digital circuit design, the NAND and NOR gates could potentially enable more sophisticated genetic circuits. The two-input NAND and NOR gates regulate *mtrC* gene expression using tailored sensing blocks that respond to common small molecule inputs - IPTG and OC6 for the NAND gate, or OC6 and aTc [aTc = anhydrotetracycline] for the NOR gate (Extended Data Figure 4a and 4b). Plasmids encoding the NAND (pNAND-*mtrC*) and NOR gates (pNOR-*mtrC*) gates were transformed into △*mtrC* mutants and strains were inoculated into OECT devices. Following overnight inoculation, channel currents were measured, and the corresponding decay rate constants were evaluated and plotted as 2D heat maps to present the response to varying inducer concentrations (Figure 4a and 4b). As expected, the NAND and NOR gates both yielded current-decay rate constants conforming to the expected truth tables, demonstrating the direct conversion of transcriptional logic to electrical signals by the OECT. Additionally, we noticed similar decay profiles of the channel current I_DS_ in mutants expressing logic ’1’ in the NAND and NOR gates. Conversely, the profile for the logic ’0’ gate samples bore resemblance to that of negative controls lacking *mtrC* expression (Extended Data Figure 4e and 4f). Together, these results demonstrate that transcriptional logic can be coupled to an OECT output.

**Figure 4.**
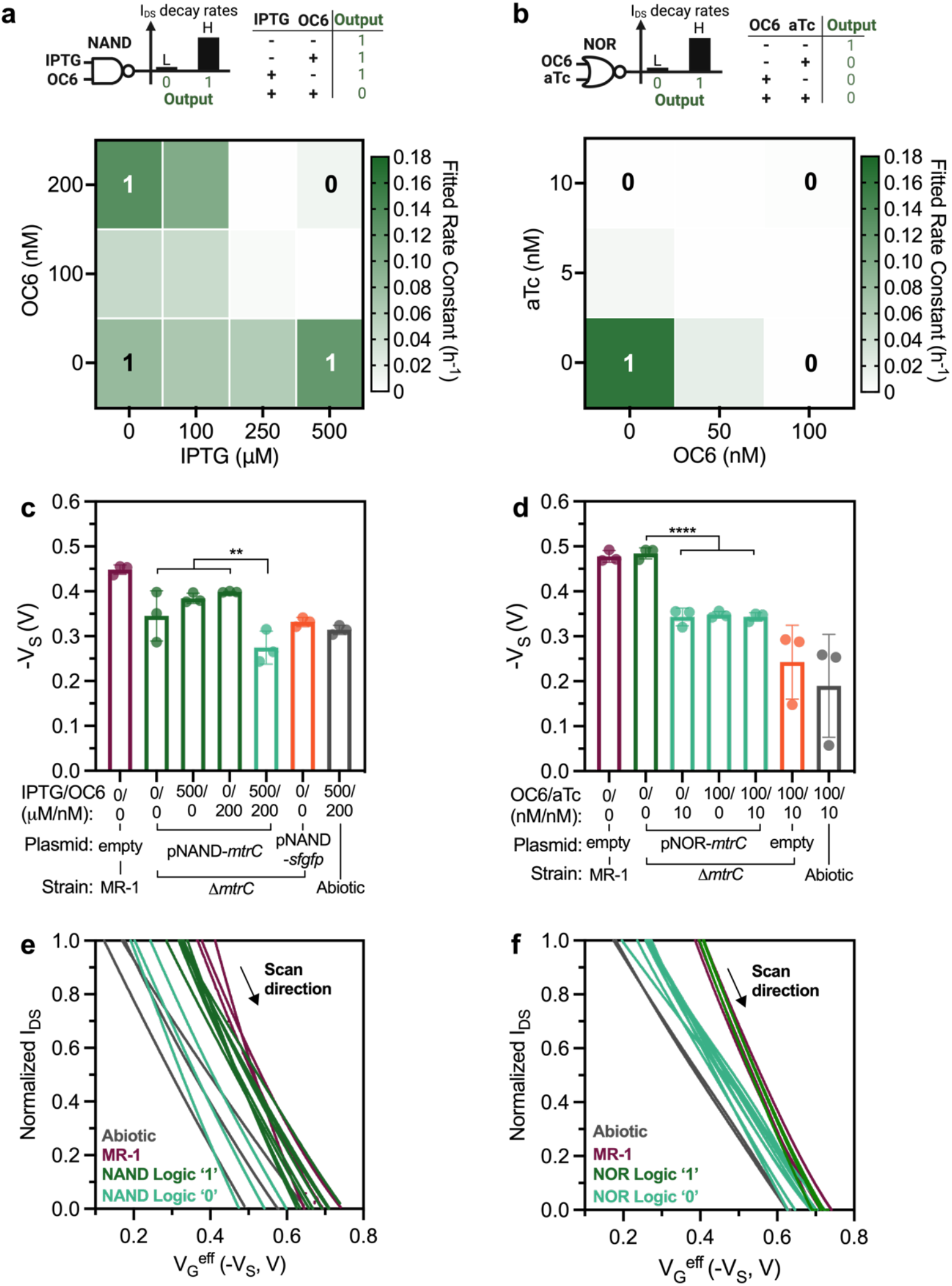
The OECT responds to strains carrying genetic Boolean logic gates. I_DS_ decay rate constants plotted with combinatorial inducer concentrations of △*mtrC* mutants carrying (**a**) NAND and (**b**) NOR Boolean logic gates. Measured source potentials V_S_ were extracted at V_GS_ = 0 V for the (**c**) NAND and (**d**) NOR gate samples. Channel current I_DS_ were normalized to the range in the transfer curves for (**e**) NAND and (**f**) NOR gate samples. Representation of p-values (n.s. p > 0.05, * p ≤ 0.05, ** p ≤ 0.01, *** p ≤ 0.001, **** p ≤ 0.0001) and data show the mean ± SD of 3 biological replicates.

Similar to the de-doping mechanism investigation, we used an Ag/AgCl pellet RE to monitor the electrode potential with the negative of the measured source potential used as the effective gate potential, V_G_^eff^ = -V_S_. To ensure complete turn-on and turn-off behavior, mutants were induced using a combination of maximum and minimum concentrations of inducers, corresponding to the corners of the respective 2D heat map (Figure 4a and 4b). For instance, 100 nM of OC6 and 10 nM of aTc were used for the NOR gate. The source potentials were measured 24 hours after inoculation against Ag/AgCl RE with the applied V_GS_ = 0 V for all induced mutants and controls. As depicted in Figure 4c and 4d, the mean source potential decreased by 101.4 mV and 139.6 mV for logic 1s compared to logic 0s for NAND and NOR gates, respectively. The shifts in source potential are clear evidence for reduction of the source electrode and channel by mutants successfully controlling EET flux according to the predicted circuit logic. Channel reduction in response to *mtrC* expression was further supported by transfer curves. To mitigate variations in transfer curve slopes resulting from channel thickness or fabrication inconsistencies (Extended Data Figure 4c and 4d), I_DS_ values were normalized to the range of 1. As shown in Figure 4e and 4f, we measured a clear shift toward more negative V_S_ (or positive V_G_^eff^) for mutants expressing logic 1s (NAND or NOR gates were on) and positive *S. oneidensis* MR-1 controls. Leaky gene expression and minimal EET activities from ‘off’ mutants could be the cause of the slight overall shift in the respective V_S_ and transfer curves. Overall, the integration of transcriptional logic with OECTs creates a general and streamlined platform that unlocks the diverse computational power available to biological systems.

### Synaptic behavior in OECTs containing electroactive bacteria

In addition to sensing applications, OECTs have emerged as promising platforms for emulating synaptic plasticity due to their analogous working principle as synapses. In a biological synapse, the presynaptic action potential releases neurotransmitters across the synaptic cleft that modulate the postsynaptic membrane potential^42^. When operating as a neuromorphic element, an OECT uses the gate voltage as the presynaptic input and the channel current as the postsynaptic output (Figure 5a). Because the channel current is a function of ion diffusion as well as redox processes within the electrolyte, the history of presynaptic inputs can influence channel conductance, resulting in a memory effect within the device. Thus, we explored whether *S. oneidensis* MR-1 could alter synaptic weight in an OECT in a genetically controlled manner.

**Figure 5.**
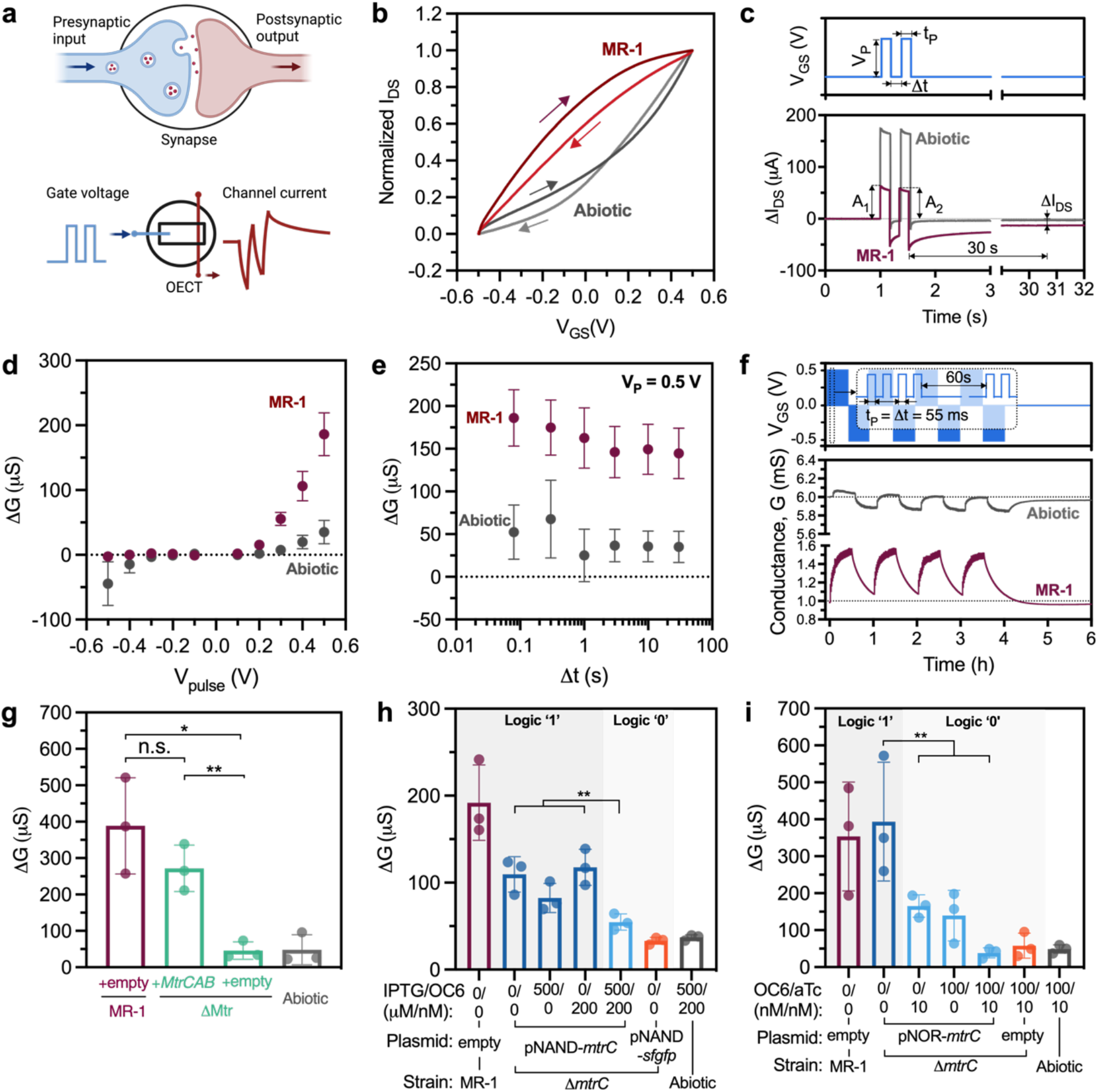
Synaptic behaviors of OECTs inoculated with electroactive bacteria. (**a**) Analogous to the neuronal synapse, the OECT employs gate voltage as the presynaptic input and channel current as the postsynaptic output. (**b**) Transfer curves of OECTs inoculated with *S. oneidensis* MR-1. (**c**) Illustrations of the paired pulse input from the gate (c, upper plot), and the corresponding channel current changes △I_DS_ of biotic and abiotic OECTs (c, lower plot). (**d**) The channel conductance changes △G plotted with varying V_P_, while t_P_ = △t = 80 ms. (**e**) Channel conductance changes △G for varying pulse interval △t, with fixed t_P_ = 80 ms, V_DS_ = −0.05 V, and V_P_ = 0.5 V. (**f**) Continuous voltage pulses were applied to the gates (f, upper plot) and the corresponding channel conductance for *S. oneidensis* MR-1 inoculated OECTs (f, lower plot). (**g**) Conductance changes △G for OECTs inoculated with △Mtr knockout strains complemented with either *MtrCAB* Buffer gate or empty vector plasmids. The △G corresponds to △*mtrC* mutants carrying (**h**) NAND and (**i**) NOR Boolean logic gates with different inducer combinations. Representation of p-values (n.s. p > 0.05, * p ≤ 0.05, ** p ≤ 0.01, *** p ≤ 0.001, **** p ≤ 0.0001) and data show the mean ± SD of 3 biological replicates.

A crucial property of synaptic transistors is non-volatility, which can be visualized as hysteresis in the transfer curve. At a channel voltage of V_DS_ = −0.05 V, we measured transfer curves of OECTs containing *S. oneidensis* cells by cycling the gate voltage between −0.5 V and 0.5 V. As shown in Figure 5b, *S. oneidensis* MR-1 inoculated OECTs showed a marked hysteresis profile relative to abiotic controls, suggesting a form of memory endowed by the bacteria. Encourage by this result, we evaluated the synaptic behavior of these devices. Two forms of short-term plasticity (STP), namely, paired pulse facilitation (PPF) and paired pulse depression (PPD) are commonly examined for their importance in decoding temporal information. As shown in Figure 5c, the paired pulses were defined by the pulse amplitude V_P_, pulse duration t_P_, and pulse interval △t. The PPF and PPD indices were defined as A_2_/A_1_ * 100%, where the A_1_ and A_2_ represent the channel current pulse amplitude relative to the pre-pulse level, induced by the paired gate voltage pulses. For simplicity, the A_2_/A_1_ index will be used instead of PPF and PPD in the following sections. The synaptic weight was defined as channel conductance (G) and the weight change was measured 30s after the end of the last gate voltage pulse.

To compare the effects of pulsed gate voltages (V_P_) on the channel responses and associated short-term memory behavior, the pulse duration t_P_ and pulse interval △t were initially kept constant at 80 ms. As shown in Figure 5d, abiotic OECTs displayed small and symmetrical weight changes to the varying gate pulse voltages. On the other hand, *S. oneidensis* MR-1 inoculated OECTs exhibited minimal conductance changes to negative gate pulses, while the conductance increased sharply with more positive V_P_. The same asymmetric response to positive versus negative pulses was also observed in the A_2_/A_1_ index, where only a marked decrease was observed with more positive V_P_ from biotic OECTs (Extended Data Figure 5a). Next, we tested the spike timing dependence by varying the gate pulse interval (△t) while maintaining a constant pulse duration t_P_ of 80 ms and pulse voltage V_P_ of 0.5 V or −0.5 V. Interestingly, for *S. oneidensis* inoculated OECTs, the channel conductance increased with positive pulses across a range of spike timing (Figure 5e), while the conductance changes were negligible with negative pulses (Extended Data Figure 5b). Consistent with our previous results, minimal and symmetrical conductance changes were observed in abiotic OECTs regardless of the pulse interval △t and pulse voltage. The A_2_/A_1_ index was also measured with different spike timing. For positive pulses, the A_2_/A_1_ index for all samples decreased exponentially as △t decreased, with biotic OECTs exhibiting more pronounced changes (Extended Data Figure 5c). Conversely, when responding to negative pulses the A_2_/A_1_ index for biotic OECTs was only distinguishable from the abiotic ones when the △t decreased to 80 ms (Extended Data Figure 5d). Together, the *S. oneidensis* MR-1 inoculated OECTs exhibited distinct synaptic behaviors, responding exclusively to positive gate voltages with a short-term increase in synaptic weight. Conversely, negative pulses induced negligible synaptic response in the biotic OECTs, comparable to the responses observed in abiotic OECTs.

To examine the reversibility of the synaptic behaviors and long-term plasticity, we subjected OECTs to continuous pre-synaptic stimuli. As depicted in Figure 5f, the gate voltage was repeatedly pulsed with V_P_ alternating between 0.5 V and −0.5 V for a total of 8 sessions. In each session, a pulse set of 4 was repeated 30 times for every 60 s, with the pulse duration t_P_ and interval △t maintained at 55 ms. Conductance baselines were obtained by removing the spiking during the gate pulses for better visualization. The corresponding channel conductance for the *S. oneidensis* MR-1 inoculated OECTs exhibited consistent responses to repeated stimuli sessions (Figure 5f, Extended Data Figure 5f). The conductance baselines for each continuous stimuli session were fitted with a one-phase exponential mode, while the positive session marked as S1, S3, S5, and S7 (Extended Data Figure 5g) and negative ones marked as S2, S4, S6, and S8 (Extended Data Figure 5h). The average time constants for positive and negative stimuli were 429.5 s ± 18.6 s and 1083.1 s ± 71.0 s, respectively (Extended Data Figure 5i). The small variation in the fitted time constants indicate consistent and reversible synaptic modulation of the *S. oneidensis* inoculated OECTs. In accordance with the paired pulse response, the saw-tooth channel current curves in Extended Data Figure 5g demonstrate the strong transient doping of the channel by the positive pulses and rapid de-doping during resting intervals (V_GS_ = 0V). The rapid de-doping during resting also suggests minimal long-term plasticity (LTP) induced by the presynaptic pulse in the biotic OECTs. However, further experiments on cell metabolic rates and viability are necessary to understand the long-term stability and plasticity of the biotic OECT.

Finally, after establishing the basic synaptic behaviors of OECTs containing *S. oneidensis*, we examined the correlation between EET and synaptic modulation using EET-deficient knockout strains complemented with *mtrC* or *mtrCAB*. For these experiments, paired inputs of V_P_ = 0.5 V and t_P_ = △t= 80 ms were used. EET-deficient mutant strains (△*mtrC* and △Mtr) containing the appropriate plasmids were inoculated into OECTs following steady state gene expression. As expected, strains with reconstituted Mtr pathways (△*mtrC* +*mtrC* and △Mtr +*mtrCAB*) showed similar conductance changes and A2/A1 index values to those of *S. oneidensis* MR-1, while negative controls (△*mtrC* +empty and △Mtr +empty) behaved similarly to abiotic OECTs (Figure 5g, Extended Data Figure 5e). These data suggest that the synaptic functions in biotic OECTs are directly correlated with EET activity from the Mtr pathway. To further examine the extent of genetic control over synaptic behavior, the △*mtrC* strains carrying Boolean logic gates (NAND and NOR) were subjected to different inducer combinations. As shown in Figure 5h and 5i, strains expressing logic output 1 had marked weight changes compared to the logic output 0 strains, demonstrating computational control over synaptic function in response to specific environmental cues. Although the detailed mechanisms underlying the correlation between synaptic behavior and bacterial EET are not yet clear, our current findings establish a direct link between the programmable synaptic response and cellular EET. By demonstrating the direct involvement of EET in shaping synaptic behavior, these results pave the way for further studies on artificial synapses and biocomputing.

## Discussion

OECTs are powerful devices for interfacing with biological systems because they inherently couple ionic and electronic signaling. While the majority of OECT applications leverage biological changes to alter ion transport or induce electrochemical changes in the channel, we demonstrated that biological electron transport from living bacteria can also be coupled to OECT sensing and computation. Specifically, we found that the model electroactive bacterium *S. oneidensis* MR-1 could change the conductivity and doping state of the PEDOT:PSS channel via EET. This process was directly tied to the presence of metabolically active cells. By controlling the bias potential, the kinetics of electron transfer between cells and electrodes could be modulated, providing a means to tune electron transfer to the channel^43^. During our OECT operation, the use of a polarizable Au gate instead of a non-polarizable Ag/AgCl gate could introduce inaccuracies when measuring the channel potential due to capacitive effects on the Au gate and uncertainties in the onset of redox reactions on the gate and channel due to the lack a fixed electrode potential^44,45^. Consequently, to enable precise measurements of the source potential, we used Ag/AgCl pellet pseudo-reference electrodes (RE) to investigate the de-doping mechanisms. Combined with UV-Vis spectroscopy and alternative OECT designs, we showed that *S. oneidensis* can interact with both the gate and the channel, resulting in charge accumulation and change in the doping state of the channel. Furthermore, we established that the gate bias voltages could be leveraged to tune electrode potential, consequently regulating EET to the electrode and channel. This gate-voltage-controlled modulation of cellular EET de-doping efficacy underscores the intricate interplay between biological and electronic components in the hybrid OECTs. However, the electron transfer and the de-doping via the cell-channel interface (path 1 in Figure 2b) can still mingle with the cell-gate path unless extreme bias potentials are used, which could result in undesirable background electrochemical reactions and stress on bacteria cells. Thus, when investigating gate- induced doping with EET (path 2 in Figure 2b), it may be advantageous to only allow EET between the cell and gate electrode while segregating the channel from any EET activities. Although we did not isolate bacteria to one part of the device, future OECTs could accomplish this using light-patternable biofilms^46^, ion-permeable membranes to separate the gate and channel, or multiple gates^47^. Alternative device architectures, such as the use of floating gates, could also be employed to enhance amplification from EET^45,48^.

On the timescale of our experiments, direct EET through the Mtr pathway dominated, but flavins also accelerated channel de-doping when added exogenously. Biofilm formation was also not required to observe the desired response result, consistent with previous studies on OECTs containing *S. oneidensis*^18^. This is promising because genetic circuits typically function best in planktonic, actively growing bacteria^49^. The diminished role of flavins during normal growth in our OECTs is in contrast to previous studies of *S. oneidensis*-PEDOT:PSS composite electrodes, which found that flavins were critical for current generation^50^. These differences can likely be explained by different surface morphology, hydrophobicity, and other chemical factors affecting the conducting polymer in the bacteria-PEDOT:PSS composite electrode, as compared to the PEDOT:PSS film utilized in our work. Indeed, post-processing of PEDOT:PSS can drastically alter its conductivity, surface properties, and redox potential^51^; these factors likely dictate polymer-bacteria interactions and warrant more systematic investigation. Finally, a variety of other (semi)conducting polymers have been evaluated in OECTs. Future research aimed at exploring the interaction of *S. oneidensis* and new materials^52^, especially n-type conductive polymers, is likely to result in OECTs with enhanced sensitivity, response time, operation efficiency, and other desirable characteristics.

Relative to traditional microelectronics, living systems offer several computational advantages such as enhanced efficiency, self-repair, and the inherent capability for parallel, distributed, and adaptive computation. These features make living systems well-suited for various applications, including biosensing and novel computational paradigms like neuromorphic computing, amorphous computation, and morphological computation^53^. To showcase the potential advantages of electroactive bacteria-inoculated OECTs in biosensing and biocomputing, we utilized plasmid-based Boolean logic gates to control cellular EET flux, achieving the direct conversion of transcriptional logic into electrical signals in response to combinations of chemical stimuli. Our modular genetic circuits regulated the expression of different parts of the Mtr pathway, since we demonstrated that these proteins directly contribute to changes in channel conductivity. However, enzymes and proteins upstream of this pathway in *S. oneidensis* or other Mtr homolog-containing bacteria (*Aeromonas hydrophillia*, *Vibrio natrigens*, etc.) may be enticing targets for future engineering since metabolic flux is directly connected to EET^54,55^. We chose transcriptional regulation for our circuits, but in principle any type of genetic regulation could be connected to an OECT output. For example, ferredoxin circuits^24^, integrases^14,56^, anti-repressors^57^, and other genetic regulatory motifs could be used to optimize signal transduction, dynamic range, and gate simplicity^58^. Overall, our system demonstrates the translation of Boolean logical computations in *S. oneidensis* to electrical readouts. Within this EET framework, we envision OECTs as a foundational platform for connecting practically any metabolic, genetic, or protein-based circuit to an electronic output.

Lastly, as a demonstration of future biocomputing applications, we employed OECTs containing electroactive bacteria as a platform to emulate synaptic behavior. Leveraging inducible control over EET flux, we established a correlation between EET and synaptic modulation. Biotic OECTs exhibited distinct paired pulse responses and conductance changes compared to abiotic devices. With positive gate pulses, the channel exhibited a notable spike-recovery response to both the rising and falling edge of the gate pulse, resulting in a short-term increase in the conductance and doping state of the channel. The spike-recovery response is likely due to the relatively low V_DS_ (−0.05 V) compared to V_GS_ (up to 0.5 V) where the transport of ions from the electrolyte into the channel outpaces the hole extraction rate within the channel^8^. However, when applying negative gate pulses, the channel exhibited a step-like response and little short-term conductance change. Although the cause of the observed asymmetric modulation and spike-recovery behavior remains unclear, comparing the results between strains with different EET protein expression levels, it is evident that the channel response arose from cellular EET, and cannot be explained by the blockade of ionic movement from the presence of cells alone. Figure S5a illustrates a distinct behavior observed in the drain electrode potential at the onset of positive gate pulse (V_GS_ = 0.5 V), wherein a sharp spike occurred below −0.9 V (vs Ag/AgCl). Conversely, as shown in Figure S5b, during the negative gate pulses (V_GS_ = −0.5 V) the drain potential increased beyond −0.3 V (vs Ag/AgCl). Given that the drain-source voltage was consistently maintained at −0.05 V, it is reasonable to assume that the channel experienced a similar potential range during the gate pulses. Consequently, we hypothesize that the channel’s low potential around −0.9 V might impede or disrupt electron transfer between the channel and attached cells, resulting in the observed asymmetric spike-recovery response to positive gate pulses. The long-term plasticity of the biotic OECTs is mainly dependent on cellular EET, as cells de-dope the channel, the conductance decreases accordingly. Further work is needed to unravel the mechanism of EET incurred spike-recovery channel response and improve the practicality of the biotic OECTs as artificial synapses. For example, reference electrodes can be used to apply a bias voltage to the channel, allowing the channel potential to be precisely controlled or monitored to study its dynamics. Additionally, these reference or auxiliary electrodes offer a convenient means of resetting the OECT in artificial synapse applications by independently controlling the doping state of the channel. New channels utilizing n-type or inorganic semiconducting materials could also provide valuable insights into the redox behaviors of the electroactive bacteria and expand device designs to enable more efficient synapses or complementary OECTs^13,59^. Similarly, new biopolymers and cell-derived materials, such as light-sensitive electronic-protonic conductors^60^, microbial nanowires^61^, and cell-secreted dopamine^62^ portend an ongoing fusion of materials, living cells, and electronics. Ultimately, the ability of EET to change synaptic weight in response to genetic memory, transcriptional regulation, and other forms of biological computation will enable new forms of metaplasticity in OECTs^63^.

OECTs have emerged as an ideal platform for combining biological and electrical signaling. Despite the more accessible experimental tools for rapid genetic and metabolic manipulation of bacteria, the existing body of research connecting bacterial cellular processes with OECTs remains limited. Our study demonstrates the viability of employing OECTs to translate bacterial computation that modulates the extracellular redox environment via extracellular electron transfer (EET). We note that in additional to the transcriptional Boolean logic gates, analog biocomputing schemes are also compatible with the OECT platform and associated future system designs. The inclusion of living cells in our hybrid devices imparts unique bio-mimetic properties such as self-regulation, biological sensing and computation, and short-term synaptic memory. Overall, our work integrates knowledge and techniques from bioelectronics, synthetic biology, and electrochemistry to create a versatile platform for future biosensing and biocomputing systems.

## Supporting information

Vgs_pulse_pair_MATLAB

IDS_VGS_Pul_long_MATLAB

## Acknowledgements

Base plasmids for the NAND circuit were generously provided by the Voigt Lab via Addgene (#49375, #49376, #49377). This research was financially supported by the Welch Foundation (Grant F-1929, B.K.K.), the National Institutes of Health under award number R35GM133640 (B.K.K.), an NSF CAREER award (1944334, B.K.K.), and the Air Force Office of Scientific Research under award number FA9550-20-1-0088 (B.K.K.). A.J.G. was supported through a National Science Foundation Graduate Research Fellowships (Program Award No. DGE-1610403). AFM experiments were performed on an instrument obtained through an AFOSR DURIP award (FA9550-21-1-0148). The authors acknowledge use of shared research facilities supported in part by the Texas Materials Institute, the Center for Dynamics and Control of Materials: an NSF MRSEC (DMR-1720595), and the NSF National Nanotechnology Coordinated Infrastructure (ECCS-1542159). We gratefully acknowledge the use of facilities within the core microscopy lab of the Institute for Cellular and Molecular Biology, University of Texas at Austin. Cartoon illustrations were created using BioRender.com.

## Author Contributions

Y.G. and B.K.K. conceived the project and designed research. Y.G. performed the majority of experiments and analysis with device fabrication assistance from Y.Z., B.T., and A.D. X.J. and J.R. assisted with device characterization and analysis. A.J.G., C.M.D., I.E.M.M., and B.M.T. constructed and characterized genetic circuits. G.P. performed the statistical analysis. Y.G. and B.K.K. wrote the manuscript with input and assistance from all authors.

## Materials & Correspondence

Request for materials and correspondence should be addressed to B.K.K.

## Data Availability

Experimental data supporting the findings of this study are available through the Texas Data Repository (doi:XXX).

---

**Extended Data Figure 1.**
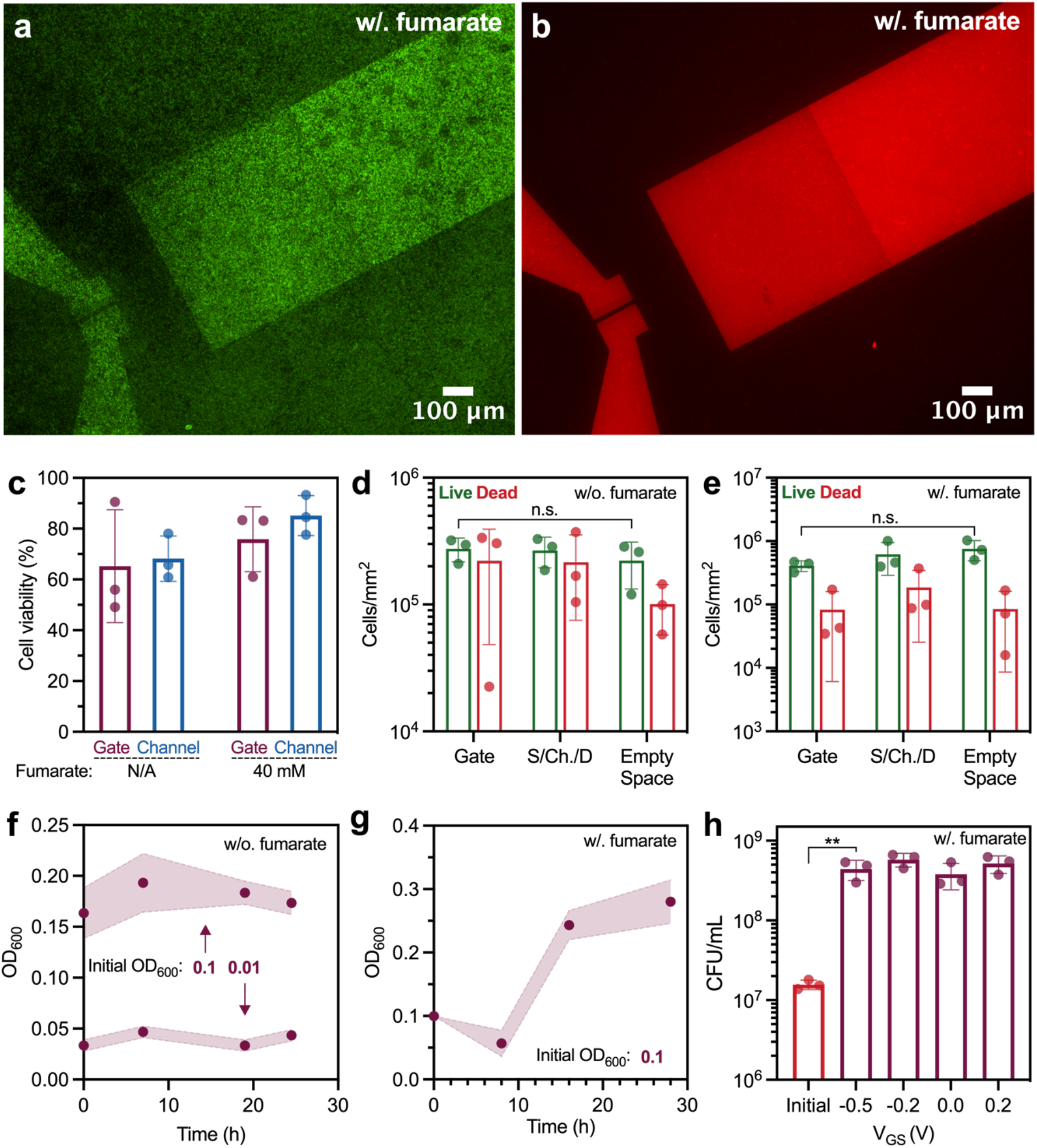
Fluorescent microscopy images showing OECT operated with constant V_DS_ = −0.05V and V_GS_ = 0.2V. Cells were supplemented with 20 mM lactate and 40 mM fumarate. LIVE/DEAD^®^ BacLight™ cell assay showing (**a**) live cells in green and (**b**) dead cells in red. (**c**) Cell viability derived from the fluorescent microscopy images. (**d, e**) Cell counts over OECT gates, source/channel/drain region (denoted as S/Ch./D), and spaces elsewhere (denoted as empty spaces) for cells supplemented with 20 mM lactate, and (**d**) without or (**e**) with 40 mM fumarate. (**f, g**) Optical Density at 600 nm (OD_600_) was measured from OECTs biased at V_DS_ = −0.05V and V_GS_ = 0.2V, cells supplemented with 20 mM lactate, and (**f**) without or (**g**) with 40 mM fumarate. The shaded region indicates the range of standard deviation. (**h**) Colony forming units (CFUs) were counted 24 hours after OECTs operation with constant V_DS_ = −0.05 V and V_GS_ biased at −0.5 V, −0.2 V, 0.0 V, or 0.2 V. Cells were supplemented with 20 mM lactate and 40 mM fumarate. Representation of p-values (n.s. p > 0.05, * p ≤ 0.05, ** p ≤ 0.01, *** p ≤ 0.001, **** p ≤ 0.0001) and data show the mean ± SD of 3 biological replicates.

**Extended Data Figure 2.**
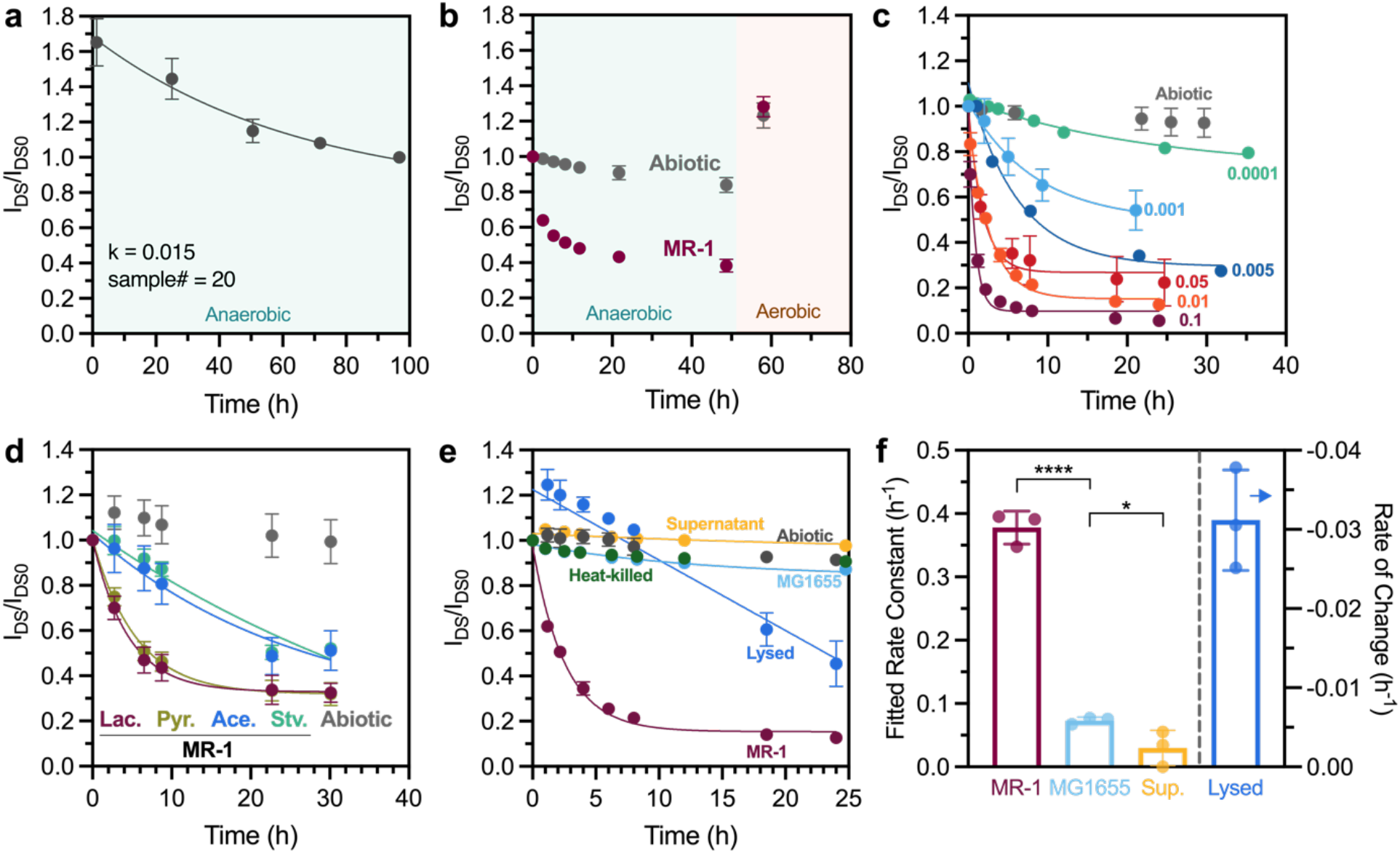
OECT channel currents I_DS_ plotted as I_DS_/I_DS0_ during (**a**) anaerobic stabilization without bacteria cells and (**b**) 48-hour operation followed by exposure to the oxygen in the ambient environment. The I_DS_/I_DS0_ curves of *S. oneidensis* MR-1 inoculated OECTs with (**c**) varying inoculation OD_600_ numbers or (**d**) supplemented with either 20 mM sodium lactate (Lac.), 20 mM sodium pyruvate (Pyr.), or 20 mM sodium acetate (Ace.) as the electron donors. No carbon source was supplied to the starved cells (Stv.) (**e**) The I_DS_/I_DS0_ curves for living, lysed, and heat-killed *S. oneidensis* MR-1 cells, as well as living *E. coli* MG1655 cells. (**f**) The I_DS_ decay rate constants for living *S. oneidensis* and *E. coli* cells, as well as *S. oneidensis* supernatant. The lysed *S. oneidensis* curves were fitted using a linear regression model. Cells were supplemented with 20 mM lactate for *S. oneidensis* MR-1 and 20 mM glucose for *E. coli* MG1655, no electron acceptor was added. Initial inocula were adjusted to OD_600_ of 0.01. Representation of p-values (n.s. p > 0.05, * p ≤ 0.05, ** p ≤ 0.01, *** p ≤ 0.001, **** p ≤ 0.0001) and data show the mean ± SD of 3 biological replicates.

**Extended Data Figure 3.**
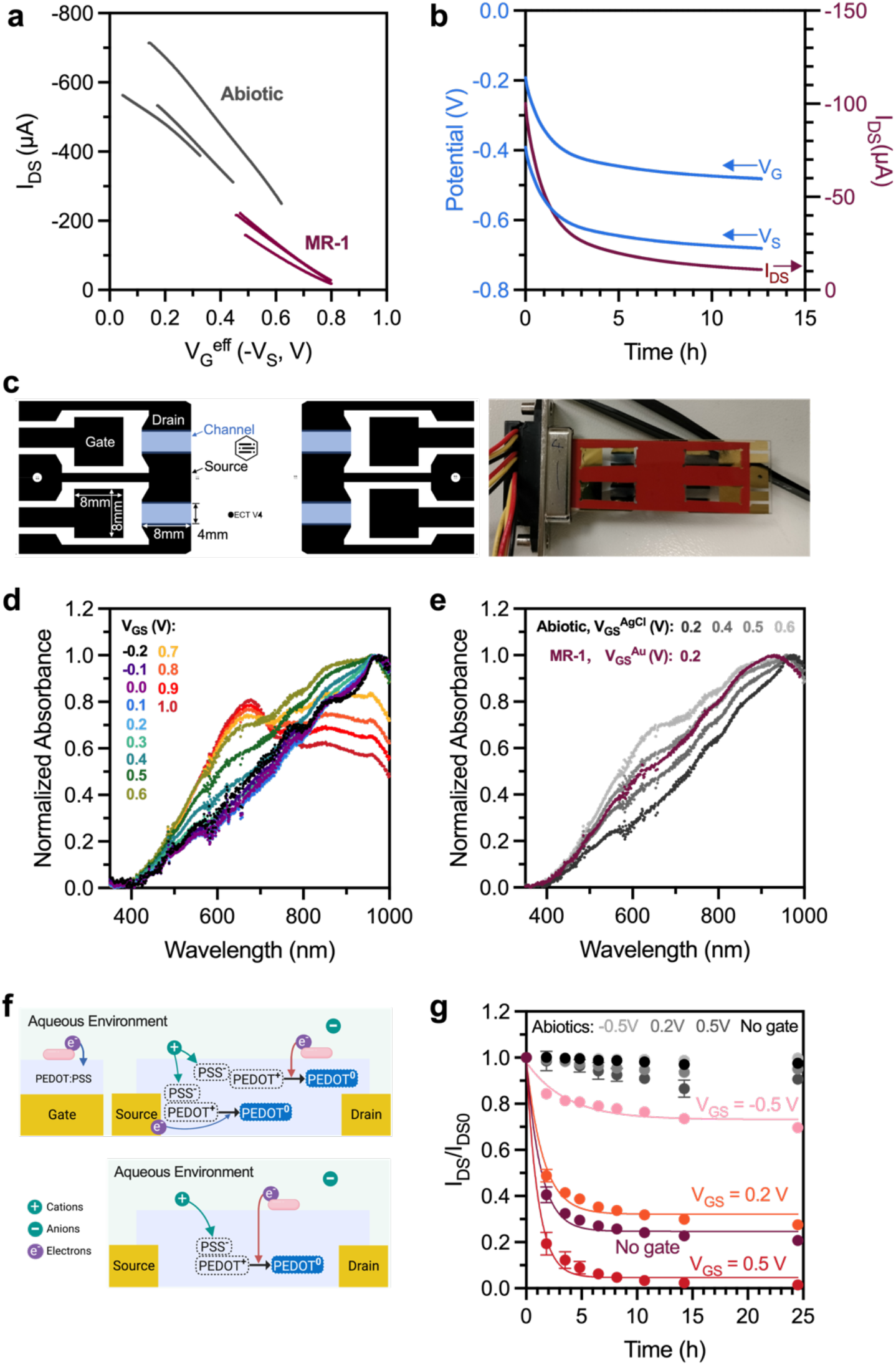
(**a**) Transfer curves plotted using negative measured V_S_ values with respect to Ag/AgCl pellet reference electrodes. (**b**) Gate and source potentials with respect to the Ag/AgCl pellet reference electrode plotted with the channel current for an OECT inoculated with *S. oneidensis* MR-1. (**c**, left) Schematic and (**c**, right) photo-image of the large channel OECT used exclusively for the UV-Vis instrument. UV-Vis spectra were collected for (**d**) abiotic PEDOT:PSS channel under different Ag/AgCl pellet gate bias voltages, and (**e**) for *S. oneidensis* MR-1 inoculated channel overlayed with abiotic channels. (**f**) Cartoon illustrations comparing the cellular EET de-doping process of OECT and the 2-electrode device. (**g**) The I_DS_/I_DS0_ curves of the 2-electrode devices and OECTs under constant gate bias conditions. *S. oneidensis* MR-1 was used for inoculum with an inoculation OD_600_ of 0.01. Data in panel (g) show the mean ± SD of 3 biological replicates.

**Extended Data Figure 4.**
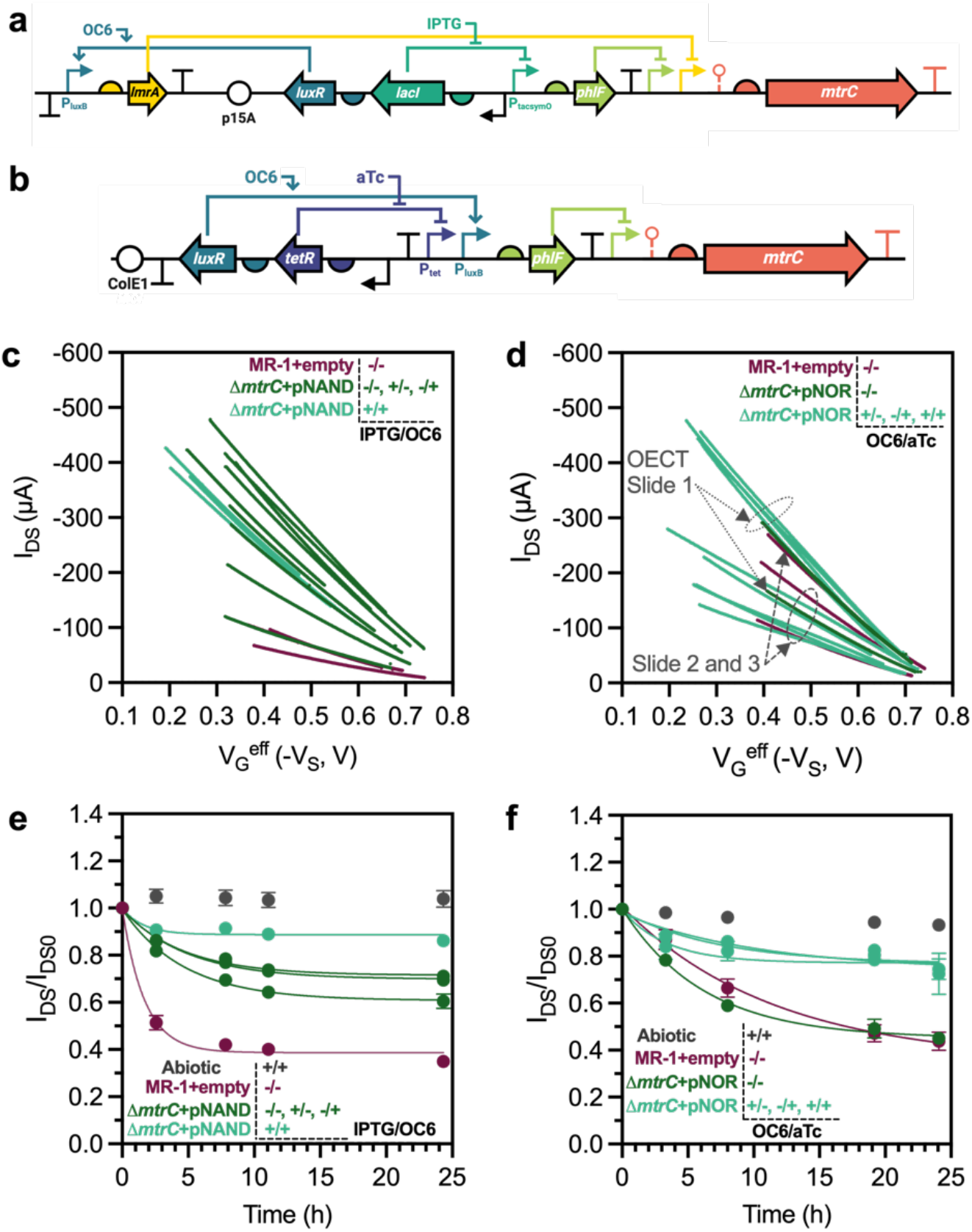
Cartoon illustrations of the plasmid architecture of the (**a**) NAND and (**b**) NOR Boolean gates expressing *mtrC*. The △*mtrC* mutants carrying the corresponding Boolean gate plasmids were brought to steady-state MtrC expression with combinations of 500 μM IPTG, 200 nM OC6, and 10 nM aTc inducers. Transfer curves of the induced (**c**) NAND and (**d**) NOR gates samples. The I_DS_/I_DS0_ for induced (**e**) NAND and (**f**) NOR gates samples. Data in panels (e) and (f) show the mean ± SD of 3 biological replicates.

**Extended Data Figure 5.**
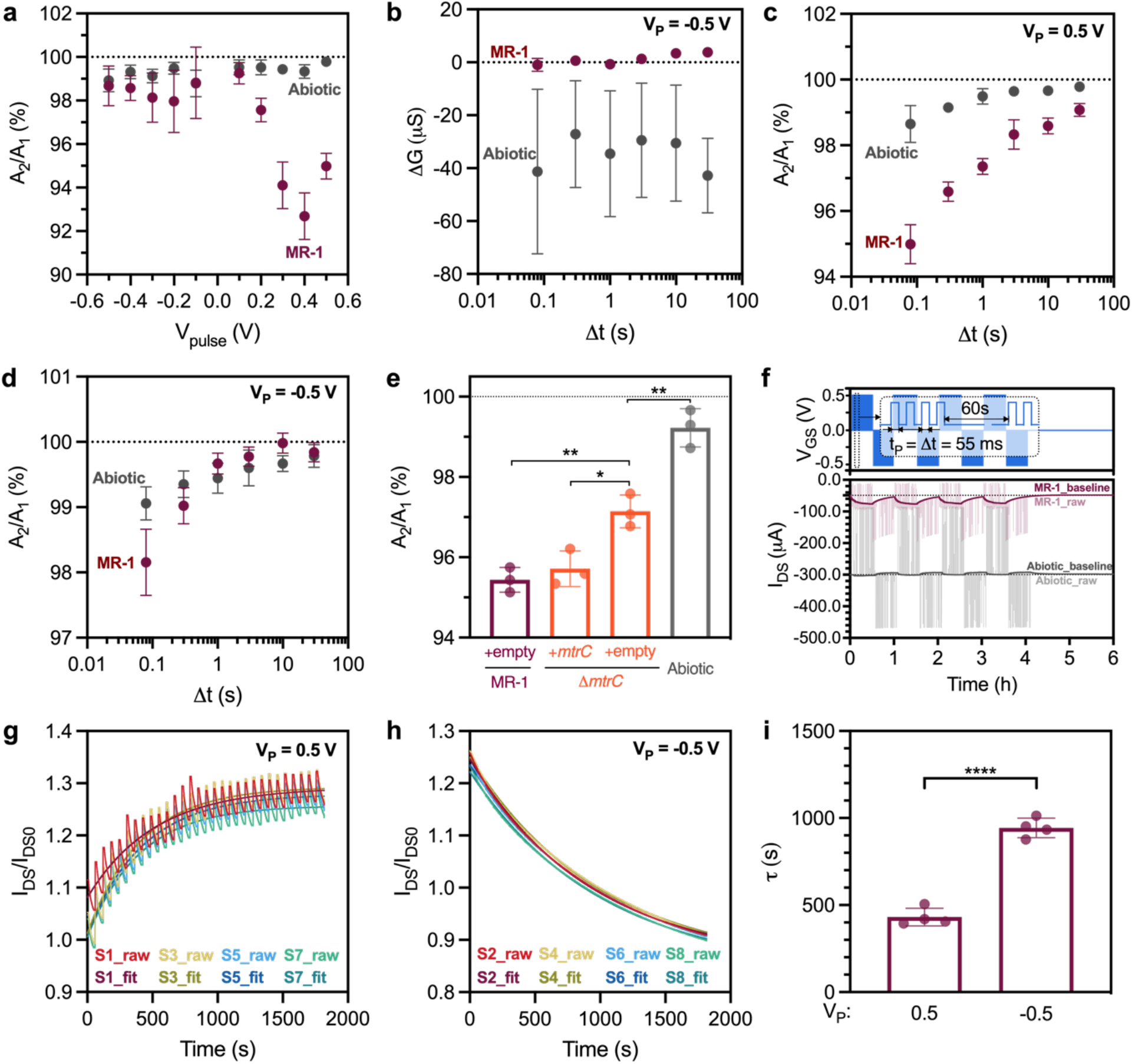
(**a**) A2/A1 index plotted with varying V_P_, while t_P_ = △t = 80 ms. (**b**) OECT channel conductance changes △G with varying pulse interval △t, fixed t_P_ = 80 ms and V_P_ = −0.5 V. A2/A1 index for OECTs with varying pulse interval △t, fixed t_P_ = 80 ms and V_P_ of (**c**) 0.5 V or (**d**) −0.5 V. (**e**) A2/A1 index of △*mtrC* strain carrying *mtrC* Buffer gate (+*mtrC*) or empty vector plasmid (+empty) under steady-state protein expression. (**f**) Channel current I_DS_ responding to the continuous voltage pulses with V_P_ of 0.5 V or −0.5V. Faded lines represent the raw I_DS_, while the bolded lines represent IDS baselines after filtering out the pulses. (**g, h**) One-phase exponential fitting of the I_DS_ baselines for each continuous 4-pulse session, with V_P_ equal to (**g**) 0.5 V or (**h**) −0.5 V. (**i**) The corresponding time constants of the fitting results. Representation of p-values (n.s. p > 0.05, * p ≤ 0.05, ** p ≤ 0.01, *** p ≤ 0.001, **** p ≤ 0.0001) and data show the mean ± SD of 3 biological replicates.

