## Supplemental Information for "A Hybrid Transistor with Transcriptionally Controlled Computation and Plasticity"

### Methods

#### *Chemicals and Reagents*

PEDOT:PSS aqueous suspension (Clevios$^{TM}$ PH1000, Heraeus Epurio LLC), ethylene glycol (anhydrous 99.8%, Sigma-Aldrich), sulfuric acid ($H_2SO_4$, 95.0-98.0 %, Sigma-Aldrich), hydrogen peroxide ($H_2O_2$, 30 wt% in water, Sigma-Aldrich), sodium DL-lactate ($NaC_3H_5O_3$, 60% in water, TCI), sodium fumarate ($Na_2C_4H_2O_4$, 98%, VWR), HEPES buffer solution ($C_8H_{18}N_2O_4S$, 1 M in water, pH = 7.3, VWR), potassium phosphate dibasic ($K_2HPO_4$, Sigma-Aldrich), potassium phosphate monobasic ($KH_2PO_4$, Sigma-Aldrich), sodium chloride (NaCl, VWR), ammonium sulfate ($(NH4)_2SO_4$, Fisher Scientific), magnesium(II) sulfate heptahydrate ($MgSO_4 \cdot 7H_2O$, VWR), Wolfe's Trace Mineral Mix (ATCC), casamino acids (VWR), isopropyl ß-D-1-thiogalactopyranoside (IPTG, Teknova), anhydrotetracycline hydrochloride (aTc, Sigma-Aldrich), 3-oxohexanoyl-homoserine lactone (OC6, Sigma-Aldrich), kanamycin sulfate ($C_{18}H_{38}N_4O_{15}S$, Growcells), Riboflavin 5′ -monophosphate sodium salt hydrate (>93 %, TCI), and LIVE/DEAD® BacLight$^{TM}$ Stain (L7012, Invitrogen), were used as received. Two-part silicone elastomer (Sylgard$^{TM}$ 184, Electron Microscopy Sciences) was used according to manufacturer instructions. All media components were autoclaved or sterilized using 0.22 µm PES filters.

#### *Bacteria Strains and Culture*

Bacterial strains and plasmids are listed in Table S1. Cell cultures were prepared from bacterial stocks stored in 20% glycerol at −80 °C. The stocks were streaked onto agar plates containing LB (for wild-type and knockout strains) or LB with 25 µg/mL kanamycin (for plasmid-harboring strains), and subsequently grown overnight at 30 °C for *Shewanella* and 37 °C for *E. coli*. Single colonies from the plates were picked and inoculated into Shewanella Basal Medium (SBM, Table S2) amended with 0.05% trace mineral supplement, 0.05% casamino acids, and supplemented 20 mM sodium lactate (2.85 µL of 60% w/w sodium lactate per 1 mL culture) for *Shewanella* and 20 mM glucose (10 µL of 2 M glucose per 1 mL culture) for *E. coli* as the electron donor. Aerobic cultures were pregrown in 15 mL culture tubes at 30 °C and 250 rpm shaking. Anaerobic cultures were pregrown using the same procedure outlined above, but with argon purged growth medium in a nitrogen-filled glovebox (S1200, Vigor) and supplemented with 40 mM sodium fumarate (40 µL of 1 M sodium fumarate per 1 mL culture) as the electron acceptor. Additional 25 µg/mL kanamycin (10 µL of 2.5 mg/mL kanamycin per 1 mL culture) was supplemented to the pregrowth medium of plasmid-harboring strains. Aerobically pregrown cultures were washed 3x using the sterile SBM growth medium and adjusted to an $OD_{600}$ of 1-3.5 (NanoDrop 2000C) before being transferred into the glovebox. For steady-state protein expression, strains were pregrown anaerobically without inducer(s) for 6 h before being diluted 1:25 into inducer-containing media (from 1000x stocks) and grown for 18-24 hours.

#### *Device Fabrication*

Three versions of the OECTs were used: small channel OECTs, 2-electrode versions of the small channel OECT without the gate, and a large channel OECT. The small channel OECTs were used for the majority of experiments with the following exceptions. In direct channel reduction experiments (Figure 2h, Extended Data Figure 3g), the 2-electrode versions of the small channel OECT were used and noted as 2-electrode devices or no gate. In UV-vis spectroscopy (Figure 2g, Extended Figure 3c – 3e), the large channel OECTs were used to fit the laser aperture of the instrument. The large channel OECTs were labeled as 'large channel OECTs' and the small channel OECTs were noted as original three-terminal OECTs or without any specific naming. OECTs were fabricated according to prior work[1]. Quartz microscopic slides (FQ-S-003, AdValue Technology) were cleaned with soapy water, acetone, and isopropyl alcohol, and dried with nitrogen before oxygen reactive-ion etching (RIE, 150 W, 50 sccm, 120 s). Quartz slides were coated with a photoresist layer (AZ5209E) and lithographically patterned to define the electrode layout. Subsequently, Au electrodes (100 nm) with a Ti adhesion layer (10 nm) were thermally evaporated on the quartz slides, and excess materials were removed with acetone lift-off. Another photolithographic pattern was formed to define the PEDOT:PSS region over the channel and the tip of the gate to obtain a width/length of 150 μm x 10 μm and 500 μm x 500 μm, respectively. The PEDOT:PSS (Clevios$^{TM}$ PH1000) was filtered (0.22 μm PES filters) and spun cast on the pattered quartz slides, followed by hot plate drying at 90 °C for 15 min and acetone lift-off. To increase the conductivity, PEDOT:PSS films were immersed in ethylene glycol at 90 °C for 3 min over a hot plate. The as-fabricated PEDOT:PSS film had an average thickness of 26.5 nm for the channel and 38.3 nm for the gate-tip layer. Polydimethylsiloxane (PDMS, Sylgard$^{TM}$ 184) with 9 % wt curing agent was drop cast and cured for 48 hours at room temperature to ensure surface smoothness. The OECT chambers and fluid access ports in the PDMS sheets were manually cut with a hole punch.

The OECTs with the larger channel size were only used for UV-vis measurement. The electrodes were fabricated in the same way as the smaller OECT. The PEDOT:PSS channel was fabricated by drop-casing 80 μL PEDOT:PSS over the entire slide, followed by 20 minutes of air-drying and heating at 90°C for 20 minutes. Afterward, the excessive PEDOT:PSS film was removed with cotton swabs soaked with 70% ethanol. Conductivity enhancement was achieved by EG treatment of 10 min at 90 °C. Silicon spacers (0.5 mm, GBL664581, Sigma-Aldrich) were hand cut to create the OECT chambers and only used with the large channel OECTs.

***Device Operation and Electrochemistry***

Shewanella Basal Medium (SBM) amended with 0.05% casamino acids and 1X Wolfe's Trace Mineral Mix was used as the base electrolyte except for the carbon source comparison experiments (Figure 2e, Extended Data Figure 2d) where the SBM was only amended with 1X Wolfe's Trace Mineral Mix. When noted, the SBM was further supplemented with 40 mM sodium fumarate to support cell growth. Before each experiment, the OECT slides and PDMS sheets were autoclaved separately and assembled in the biosafety cabinet. OECT experiments were conducted in a nitrogen-filled glovebox to create the anaerobic condition except for UV-vis spectroscopy which was conducted under ambient conditions. Media and solutions stocks were purged with argon for 15 min and stored in the glovebox. A multichannel potentiostat (MultiPalmSens4, PalmSens BV) was used for the electrochemical measurements. During continuous OECT operation, unless otherwise noted, the gate ($V_{GS}$) and drain ($V_{DS}$) voltages were biased at 0.2V and -0.05 V, respectively. For all the experiments, OECTs were stabilized in the glovebox with abiotic electrolytes and constant bias voltages for 3 days before inoculation or measurements. For OECT inoculation, cell cultures grown aerobically were initially spun down, brought into the glovebox, and diluted 10X, using the same diluents as the OECT electrolytes. Subsequently, the intermediate cultures were used to inoculate OECTs at a 1:9 ratio of cell culture to the OECT electrolyte (total 100X dilution). For instance, *S. oneidensis* supplemented with 1 µM exogenous FMN with inoculum $OD_{600}$ of 0.05 was prepared from aerobically grown cell cultures. The triple-washed cell cultures (average $OD_{600}$ at 3.38) were brought into the glovebox and diluted to the intended $OD_{600}$ of 0.5 with SBM supplemented with 1 µM exogenous FMN, creating the intermediate cultures. Then 5 µL of these intermediate cultures were inoculated into the OECTs containing 45 µL of SBM supplemented with 1 µM FMN, achieving a final inoculation $OD_{600}$ at 0.05. For steady-state expression, cells were induced and pregrown anaerobically. Strains were pregrown anaerobically without inducer(s) for 6 h before being diluted 1:25 into inducer-containing media (from 1000x stocks) and grown for 18-24 hours. Pregrown cell cultures were inoculated into the OECTs without washing at a final dilution of 100-fold.

In the carbon source experiment, SBM amended with 1X Wolfe's Trace Mineral Mix was used as the base electrolyte. Aerobically grown *S. oneidensis* MR-1 cells culture were triple-washed with SBM without a carbon source, then the cell cultures were kept at room temperature for 3 hours to induce starvation conditions[2]. Afterward, cell cultures were washed again and their $OD_{600}$ was measured. Subsequently, cell cultures were brought into the glovebox and diluted to obtain the intermediate stocks with an intended $OD_{600}$ of 0.1. The dilutions were performed with SBM supplemented with either 20 mM lactate, 20 mM pyruvate, 20 mM acetate, or no carbon source. Finally, the intermediate stocks were inoculated into OECTs containing the SBM supplemented with the respective carbon source or no carbon source at a ratio of 1:9, achieving a final inoculation $OD_{600}$ of 0.01.

In cell viability experiments, aerobically grown *S. oneidensis* MR-1 cell cultures were triple-washed with SBM supplemented with 20 mM lactate. For *E. coli* cultures, the lactate was replaced with 20 mM glucose. The densities for the washed cell cultures were measured by $OD_{600}$ and the *S. oneidensis* cultures were allocated to 3 parts: live, heat-killed, and lysed cells. Heat-killed cells were obtained by incubating at 80 °C for 20 min. Lysed cells were obtained by sonication (Qsonica 55, Qsonica LLC) for 90 s at 4 °C. The output power was set to 15 W with "ON" and "OFF" intervals of 10 s and 5 s, respectively. Then cell cultures were brought into the glovebox and diluted with SBM supplemented with 20 mM lactate to obtain the intermediate cultures at an intended OD600 of 0.1. The intermediate cultures were inoculated into the OECTs at a ratio of 1:9, achieving a final inoculation $OD_{600}$ of 0.01. The *S. oneidensis* supernatants were obtained by filtering (0.22 µm PES filters) the cell cultures (initial OD600 at 0.01) anaerobically grown in the glovebox with SBM supplemented with 20 mM lactate. The OECT electrolytes were replaced with the supernatants during inoculation.

In the electrode potential measurements with Ag/AgCl pellet reference electrode (550010, A-M Systems), the Ag/AgCl electrodes were directly inserted into the OECT chamber without salt bridges. The same Ag/AgCl pellet electrodes were used as the gate for the large channel OECTs in abiotic UV-vis measurements. Hybrid OECT experiments with electroactive bacteria were conducted between 24 - 36 hours after inoculation (initial $OD_{600}$ at 0.01). Synaptic measurements were conducted after $I_{DS}$ was stabilized for at least 1 minute at $V_{GS}$ = 0 V.

#### *Spectroscopy*

The UV-vis spectroscopy of the PEDOT:PSS channel was measured from 190 nm to 1100 nm (Agilent 8453 UV-visible Spectroscopy System). A custom sample holder was 3D printed to fit the OECT slides to the instrument. Measurements were blanked with devices lacking the PEDOT:PSS channel. When bacteria cells were present in the sample, the blank devices were likewise inoculated with the same inoculum.

#### *Fluorescence Microscopy*

Microscopy was performed using a Nikon Ti2 Eclipse inverted epifluorescence microscope. Immediately after the 24-hour operation in the glovebox, OECTs were gently washed 2x by refreshing the electrolyte with the sterile SBM supplemented with 0.05% trace mineral supplement, 0.05% casamino acids. Then, the PDMS sheets were replaced with a 0.5 mm silicon spacer to ensure the sample thickness was compatible with the working distance of the microscope. Subsequently, the OECTs were gently washed with SBM containing 0.05% trace mineral supplement, 0.05% casamino acids, and LIVE/DEAD® BacLight™ Stain mix (3 µL of SYTO 9 and propidium stocks per 1 mL) at a final solution volume of 10 µL per OECT chamber. The OECTs were then sealed with coverslips, covered with aluminum foil, and transferred out of the glovebox for microscope imaging. Bacterial counts were performed using ImageJ software.

#### *Atomic Force Microscopy*

The AFM scans were conducted using a DriveAFM (Nanosurf AG). The cantilevers (Dyn190Al) were driven with the photothermal laser (CleanDrive) under dynamic mode. OECTs were randomly selected from two fabrication batches: 3 slides of the as-fabricated OECTs (8 OECTs per slide) were used as pre-inoculation samples, and 3 cleaned OECT slides post the 48-hour inoculation served as post-inoculation samples. Two OECTs per slide were randomly chosen for AFM scans using Nanosurf CX software for data acquisition. Images for PEDOT:PSS film thickness were acquired at 90 µm x 90 µm and processed with Nanosurf CX software to correct background variations. Topology and phase images were acquired at 500 nm x 500 nm and analyzed without further process. Grain sizes were fitted by the Watershed method with the resulting histograms generated using Gwyddion software.

#### *OECT Data Processing*

Measured OECT data were processed using GraphPad Prism9 and MATLAB (R2021b update 1). The measured $I_{DS}$ data were normalized to the initial value before inoculation ($I_{DS0}$) before fitting. The $I_{DS}$ decay rate constants for all samples except the lysed *S. oneidensis* were obtained by fitting $I_{DS}/I_{DS0}$ data with an exponential decay model:

$$i_{DS}(t) = e^{k*t}$$

Where $t$ is time in hours, $k$ is the fitted $I_{DS}$ decay rate constant. Fitting was performed with the built-in one phase decay function in GraphPad Prism9. The $I_{DS}/I_{DS0}$ data for lysed *S. oneidensis* samples were fitted to a simple linear regression model:

$$i_{DS}(t) = I_{DS0} + \beta t$$

Where $t$ is time in hours, the slope $\beta$ is used as the rate of change. Fitting was performed with the built-in simple linear regression function in GraphPad Prism9.
The response function of FMN concentrations was modeled with a four-parameter logistic regression function in terms of $I_{DS}$ decay rate constants (noted here as $r$):

$$\frac{r}{R_{max}} = \frac{[S]^n}{\left(K_{\frac{1}{2}}\right)^n + [S]^n}$$

Where $n$ is the Hill coefficient, $[S]$ is the FMN concentrations, $R_{max}$ is the maximum rate constant, $K_{\frac{1}{2}}$ is the half-maximum concentration constant. Fitting was performed with the built-in four-parameter dose-response function in GraphPad Prism9.
In synaptic experiments, MATLAB scripts were used to process the measured data. The 'Vgs_pulse_pair.m' script was used to identify and extract the channel current $I_{DS}$ peak values using `islocalmax` function (Figure 5d, 5e, 5g - 5i, Extended Data Figure 5a - 5e). The 'IDS_VGS_Pul_long.m' script implemented median filtering to remove pulses and extract the baseline for channel current $I_{DS}$ responding to continuous gate pulse inputs. The filtering was achieved by using the `medfilt1` function (Figure 5f and Extended Data Figure 5f). Extracted $I_{DS}$ curves were fitted to an exponential model `b(1)*exp(b(2)*x(:,1))+b(3)` with `fitnlm` function to obtain the rate constants, where b(1), b(2), and b(3) are fitting coefficients, x is time in seconds.

All scripts will be available through the Texas Data Repository.

***Statistical Methods***

Independent samples t-tests were conducted using the built-in t-test function with the unpaired option in GraphPad Prism9.

In the 2-input Boolean gate figures (Figure 4c and 4d, Figure 5h and 5i), statistical tests were performed using R (version 4.2.3) with multicomp package (version 1.4-23). In brief, we employed a linear model fitting approach to determine the interaction term between the two inputs noted as Factor1 and Factor2, specifically lm(Results ~ Factor1 * Factor2 – 1). Then, based on the significance of the interaction terms, different linear contrasts were tested for general linear hypotheses, specifically glht(model, linfc = contrast). The significance of the interaction term was examined for p < 0.05. If the interaction term was significant, the following linear contrasts were evaluated for each specific logic gate and model parameterization:

NAND gate: $H_0$: $\mu_{++} + \frac{-1}{3}(\mu_{--} + \mu_{+-} + \mu_{-+}) = 0$

NOR gate: $H_0: \frac{-1}{3}(\mu_{++} + \mu_{+-} + \mu_{-+}) + \mu_{--} = 0$

The contrast matrices for the NAND and NOR logic gates in the general linear hypothesis test are as follows: $(^2/_3, -^2/_3, -^2/_3, 1)$ for NAND gate, and $(-^2/_3, ^2/_3, ^2/_3, 1)$ for NOR gate. These contrast matrices were calculated by algebraic substitution given the following coefficient/parameter relationships defined by our parameterization of the linear model, specifically: $\mu_{++}$ = coef[1], $\mu_{+-}$= coef[1] + coef[3], $\mu_{-+}$= coef[2], $\mu_{--}$= coef[2] + coef[3] + coef[4]. The resulting p-values and the interaction term are outlined in Table S3.

If the interaction term was not significant, a linear model without interaction was fit to the data, and the model was tested with new contrast matrix as: $(-1, ^1/_3, ^1/_3, ^1/_3)$ for the NAND gate, and $(1, -^1/_3, -^1/_3, -^1/_3)$ for the NOR gate. The p-values are outlined in Table S3 without an interaction term.

### Tables

**Table S1**. Strains and plasmids used in this study.

| Strain and Plasmid | Description | Source |
|---|---|---|
| **Strains** | | |
| *Escherichia coli* | | |
| MG1655 | Wild-type strain | Lydia Contreras, University of Texas at Austin |
| *Shewanella oneidensis* | | |
| MR-1 | MR-1 (ATCC700550) wild type strain | American-Type Culture Collection |
| △*mtrC* | JG596, deletion of genes *mtrC*, *omcA*, and *mtrF*. | Jeffrey Gralnick, University of Minnesota[3] |
| △Mtr | JG1194, deletion of genes *mtrC*, *omcA*, *mtrF*, *mtrA*, *mtrD*, *dmsE*, *SO4360*, *cctA,* and *recA*. | Jeffrey Gralnick, University of Minnesota[4] |
| △*bfe* | deletion of genes *SO0702.* | Jeffrey Gralnick, University of Minnesota[5] |
| △*lysis* | S2933, deletion of genes *SO2966* and SO2974. | Lydia Contreras, University of Texas at Austin[6] |
| **Plasmids** | | |
| pCD8 | Empty Buffer gate | Ref.[7] |

| | | |
|---|---|---|
| pCD24r1 | *mtrC* Buffer gate ($sRBS1_{mtrC}$) | Ref.[7] |
| pAT4 | *Mtr* Buffer gate | This work |
| pNAND-*mtrC* | *mtrC* NAND Boolean logic gate | Ref.[8] |
| pNAND-*sfgfp* | *sfgfp* NAND Boolean logic gate | Ref.[8] |
| pNOR-*mtrC* | *mtrC* NOR Boolean logic gate | Ref.[8] |
| pNOR-*sfgfp* | *sfgfp* NOR Boolean logic gate | Ref.[8] |

**Table S2**. Shewanella Basal Medium (SBM) formulation

| Ingredient | Quantity per 1 L |
|---|---|
| $K_2HPO_4$ | 225 mg |
| $KH_2PO_4$ | 225 mg |
| NaCl | 460 mg |
| $(NH_4)_2SO_4$ | 225 mg |
| $MgSO_4*7H_2O$ | 117 mg |
| HEPES | 100 mL of 1 M HEPES |
| Casamino acids | 5 mL of 10% casamino acids in $ddH_2O$, if needed |
| Wolfe's Mineral Mix | 5 mL of Wolfe's Mineral Mix, if needed |
| $ddH_2O$ | Adjust volume to 1 L and pH to 7.2 |

**Table S3**. Relevant statistical values

| Data Sets | pNAND | pNOR |
|---|---|---|
| **Source Potential ($V_S$)** | 2-way ANOVA interaction p= 0.00313<br><br>Contrast test p = 0.00153 | 2-way ANOVA interaction p= $1.84 \times 10^{-5}$<br><br>Contrast test p = $2.36 \times 10^{-7}$ |
| **Synaptic Conductance Change ($\triangle G$)** | 1-way ANOVA contrast test p = 0.00306 | 1-way ANOVA contrast test p = 0.00152 |

### Supplementary Figures

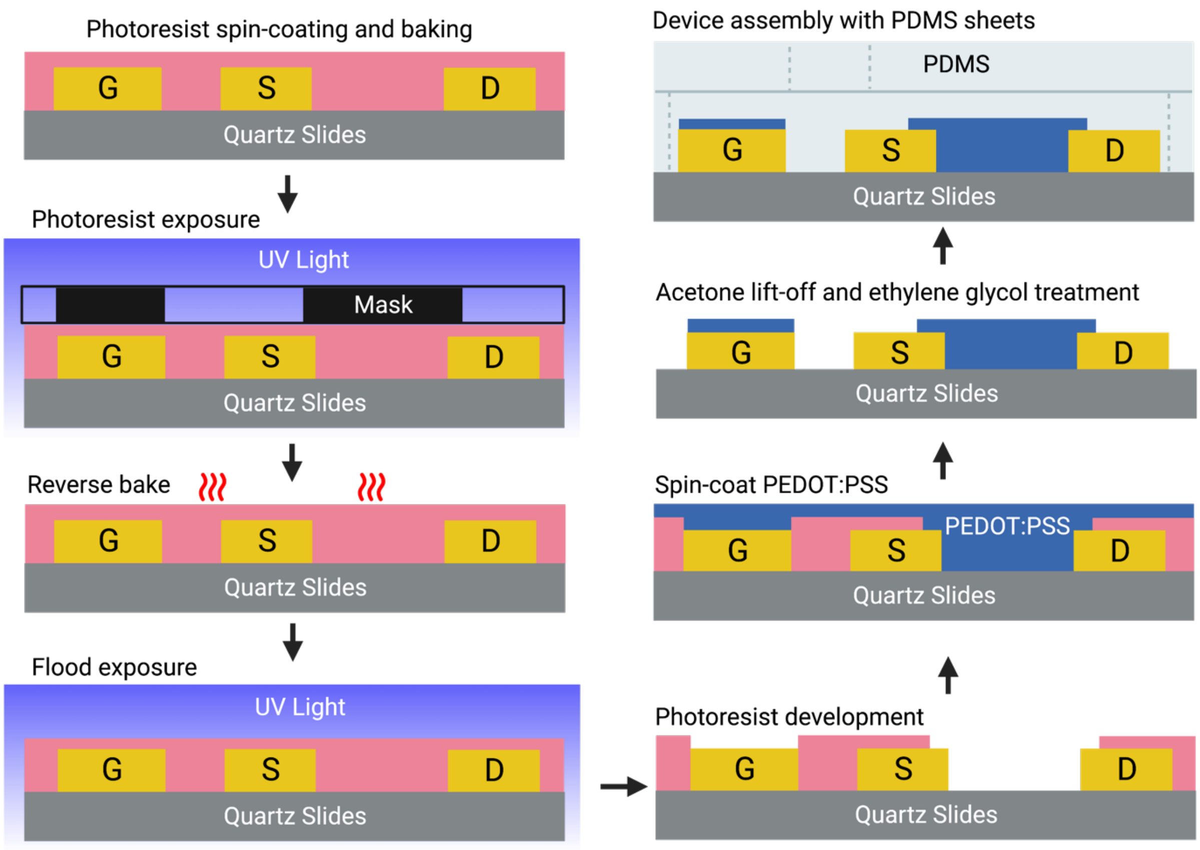

**Figure S1**. Cartoon illustration of the major OECT fabrication steps.

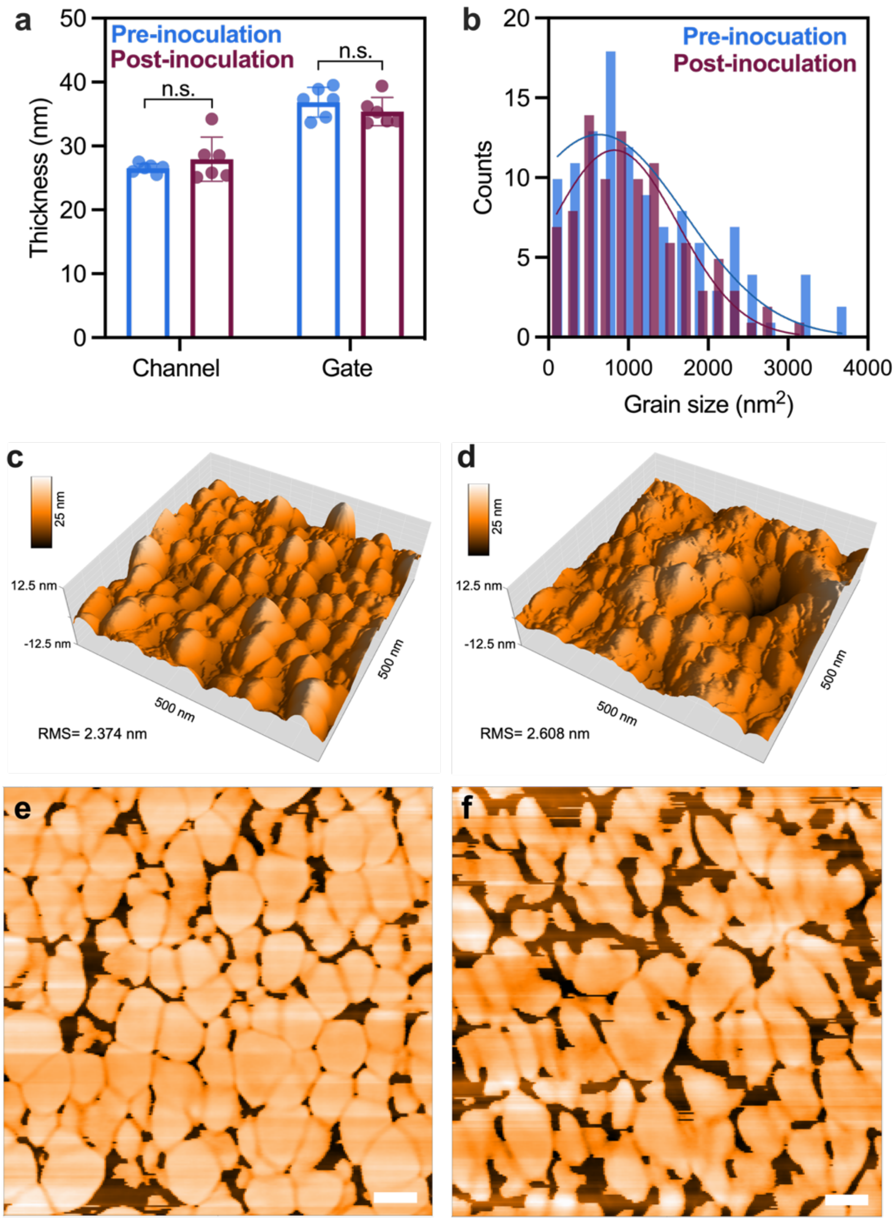

**Figure S2**. Morphological characterization of PEDOT:PSS channel and gate-tip coating. (**a**) Thickness of PEDOT:PSS films derived from AFM scans. OECTs were inoculated with *S. oneidensis* and operated with constant bias voltages of $V_{GS}$ = 0.2V and $V_{DS}$ = -0.05 V for 48 hours. (**b**) Histogram and Gaussian fit (lines) of PEDOT grain size in the channel. Surface topologies of the PEDOT:PSS channel (**c**) prior to and (**d**) after the S. oneidensis incubation. Phase images of the PEDOT:PSS channel (**e**) prior to and (**f**) after the incubation, scale bars represent 50 nm. Representation of p-values (n.s. p > 0.05, * p ≤ 0.05, ** p ≤ 0.01, *** p ≤ 0.001, **** p ≤ 0.0001) and data in panel (a) show the mean ± SD of 6 biological replicates.

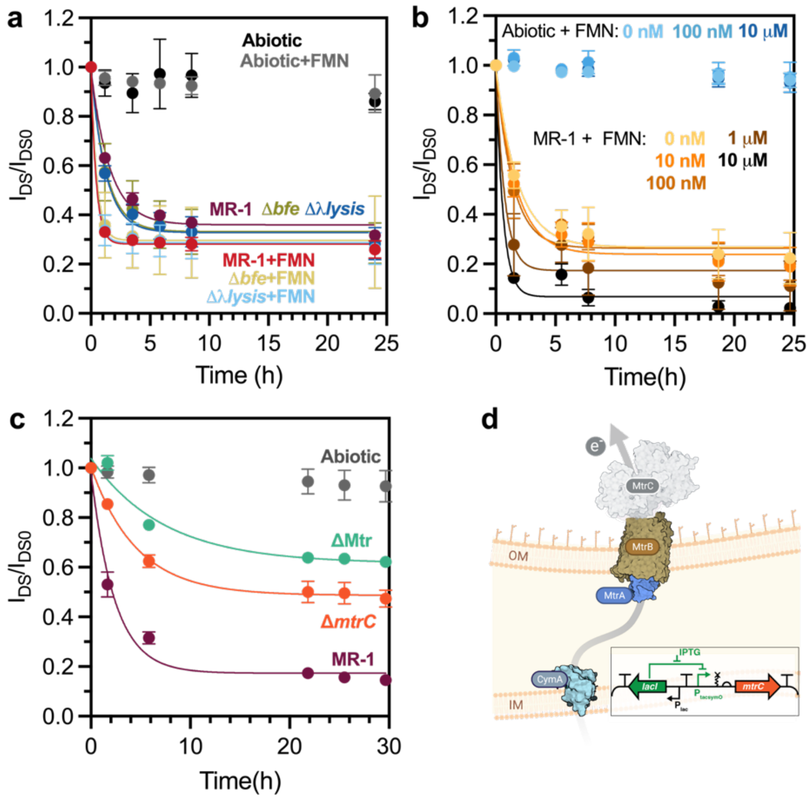

**Figure S3**. Measured OECT channel currents response to different EET mechanisms.
(**a**) The $I_{DS}/I_{DS0}$ curves of knockout strains with and without the addition of exogenous FMN (1μM), and (**b**) *S. oneidensis* MR-1 cells with varying exogenous FMN concentrations. Initial inocula were adjusted to $OD_{600}$ of 0.05. (**c**) The $I_{DS}/I_{DS0}$ curves of △*mtrC*, △Mtr, and MR-1 strains with initial inoculation $OD_{600}$ at 0.1. (**d**) Cartoon illustration of the △*mtrC* strain and (d, insert) diagram of the *mtrC* Buffer gate controlled by the IPTG inducer. Faded shapes indicate removed proteins with genomic deletion. Data show the mean ± SD of 3 biological replicates.

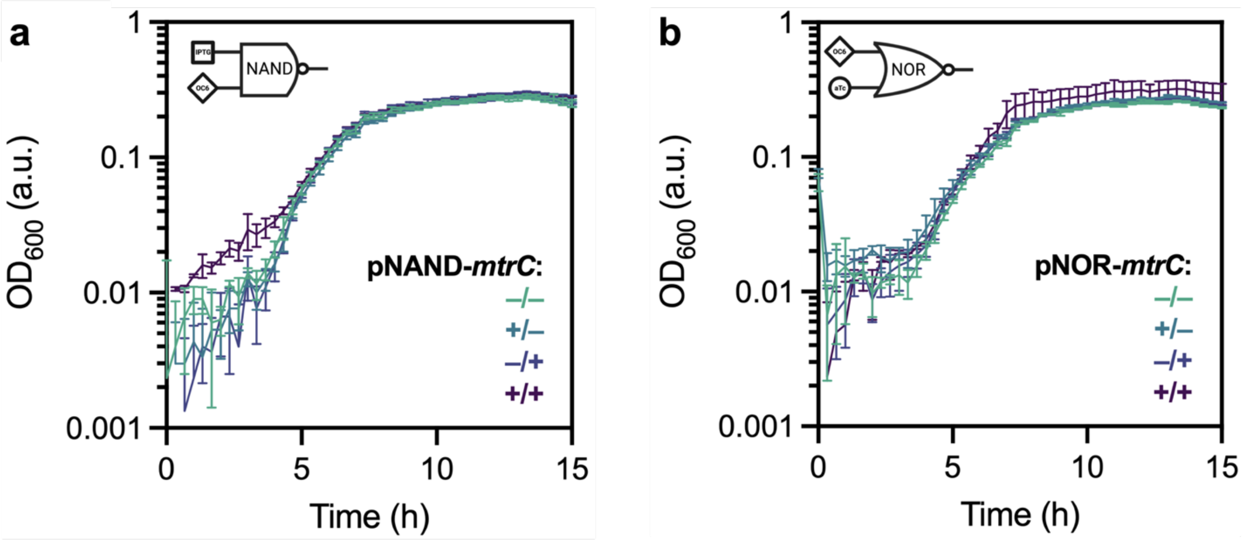

**Figure S4**. Growth curves of mutant strains carrying the Boolean logic gates plasmids. (**a**) NAND or (**b**) NOR Boolean *mtrC* plasmids under different inducer combinations. Data show the mean ± SD of 3 biological replicates.

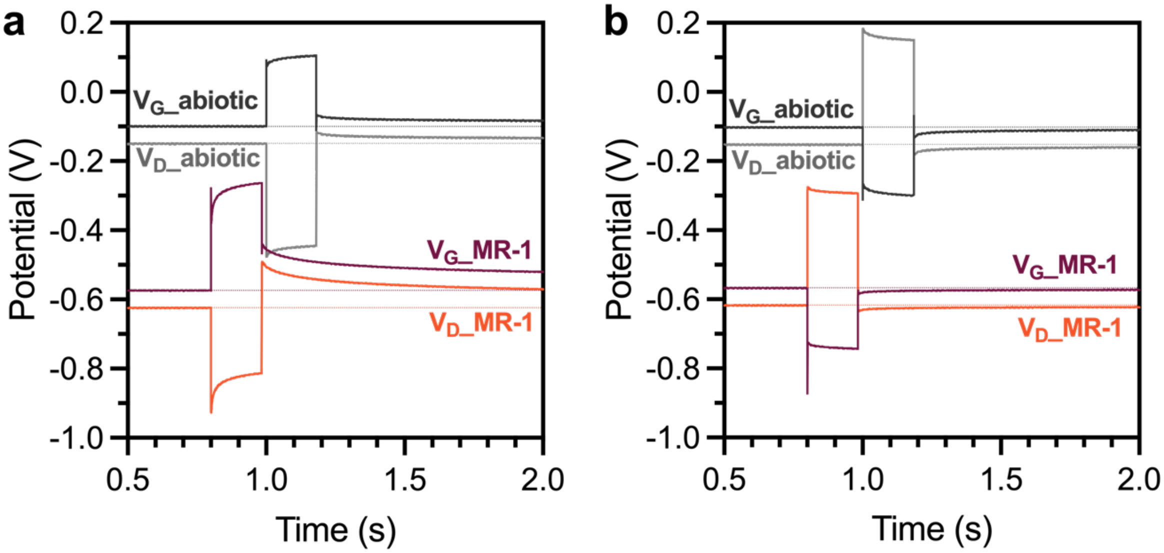

**Figure S5**. Drain and gate potentials measured against Ag/AgCl reference electrodes. Single gate pulses with $t_{pulse}$ = 1.5 s, $V_P$ equal to (**a**) 0.5 V and (**b**) -0.5 V were applied between gate and source electrodes.
