## Supplementary material for "A Hybrid Transistor with Transcriptionally Controlled Computation and Plasticity": Vgs_pulse_pair_MATLAB

% Author Yang Ga0. Keitz Group, Aug 2023
% Ch1 for channel current IDS
% IDSChx data structure: column1->time(s), column2->IDS(uA), column3->IDS
% normalized. column4->detIDS, 
% Varying pulse duration, weight change by IDS levels 30s after the 
% last VGS change
% output -> WghChgOut
%% ini For IDS measurement with VGS pulses, fast CA. 
clear;
clf;
%%%%%%%%%% adjustable variables %%%%%%%%%
IniNamStr1 = 'CH1-Abio_2pul_0V5_SBMppLKInd_JPJR.csv';
MinProminence=4;    % prominence valure for IDS pulse, may adjust if the pulse amplitude is small relativie to noise
Cutoff=2000;    % cutoff x number of data points at the end of IDS data, gate/channel syn.
PulDir=1;   % VGS pulse polarity, 1 for positive pulse (e.g. 0V to 0.5V), 0 for negative pulse (e.g. 0V to -0.5V)
WghTimAvg=0.08;    % seconds, weight averaging for initial IDS value, time before first IDS change
VDS=-0.05;      % V, channel voltage VDS, for conductance calculation
SmpInt=0.0004; % sample interval, s
WghTimDly=30;  % seconds, weight change calculation, delay time after last VGS change, 
%%%%%%%%%% end adjustable variables %%%%%%%%%
IDSIni=0;   % uA, IDS initial value before VGS pulses
IDSWgh=0;   % uA, IDS final value WghTimDly after the last VGS change
WghChgOut=zeros(10,1);   % output of weight change, change %, initial IDS, final IDS
LstVGSChgInd=0;     % IDS array index for the last VGS change
WghVGSChgInd=0;     % IDS array index for the weight calculate after time delay (WghTimDly) from the last VGS chagne
FstVGSChgInd=0;     % IDS array index for the first VGS change
WghVGSIniInd=0;     % IDS array index for the intial IDS value averaging
% read file to get data dimensions
datatableCH1 = readtable(IniNamStr1, 'ReadVariableNames', false);
IDSRawNumCH1=height(datatableCH1);
CH1Raw = zeros(IDSRawNumCH1,2); % uA, colum1->time, colum2->IDS Ch1, colum3-> IDS Ch3
IDSCh1 = zeros(IDSRawNumCH1-1-2*Cutoff,3);

%% File read

CH1Raw=table2array(datatableCH1);
% remove number of Cutoff vaules in the beginning and end of the raw data
CH1Raw(1:Cutoff,:)=[];
IDSRawNumCH1 = IDSRawNumCH1-Cutoff;
CH1Raw(IDSRawNumCH1-Cutoff:IDSRawNumCH1,:)=[];
IDSRawNumCH1 = IDSRawNumCH1-1-Cutoff;

IDSCh1(:,1)=CH1Raw(:,1);   % time, column 1
IDSCh1(:,2)=CH1Raw(:,2);   % IDS(uA), colum 2
IDSCh1(:,3)=normalize(CH1Raw(:,2),'range');   % normalized IDS to scale, column 3

%% peak detection
% get A1, A2, A1* and A2* by minimum and maximum values
TF_max=islocalmax(IDSCh1(:,2),'MinProminence',MinProminence,'MinSeparation',100);   % minSeparation 100 points, avoid noise to be considered as peak
TF_min=islocalmin(IDSCh1(:,2),'MinProminence',MinProminence,'MinSeparation',100);
% plot IDS
figure(1)
plot(IDSCh1(:,1),IDSCh1(:,2),IDSCh1(TF_max,1),IDSCh1(TF_max,2),'r*',IDSCh1(TF_min,1),IDSCh1(TF_min,2),'r+')
hold on;
%% weight change calculation
% 1. multiple peak amplitude, pulse detection
A_crs=IDSCh1(TF_min,2);   % get the A1+, A2+...
tA_crs=IDSCh1(TF_min,1);  % get the time for A1+, A2+...
Crs1Ind=find(IDSCh1(:,1)==tA_crs(1));
Crs2Ind=find(IDSCh1(:,1)==tA_crs(2));
A_crs_bas=mean(IDSCh1(Crs1Ind-2*WghTimAvg/SmpInt:Crs1Ind-WghTimAvg/SmpInt,2));

A_str=IDSCh1(TF_max,2);   % get the A1* and A2*
tA_str=IDSCh1(TF_max,1);  % get the time for A1* and A2*
Str1Ind=find(IDSCh1(:,1)==tA_str(1));
Str2Ind=find(IDSCh1(:,1)==tA_str(2));
A_str_bas=mean(IDSCh1(Str1Ind-2*WghTimAvg/SmpInt:Str1Ind-WghTimAvg/SmpInt,2));

% check if pulses on both direction were detected
if length(A_crs)==length(A_str)
    PulNum=length(A_str);
else
    if (length(A_crs)~= 0)||(length(A_str)~=0)
        if length(A_crs)>length(A_str)
            A_crs(length(A_crs),:)=[];
        else
            A_str(length(A_str),:)=[];
        end
    end
    PulNum=length(A_str);
end

% peak amplitude calculation
% for 2 or more pulse
if PulDir   % positive VGS
    d_tA_str=tA_str(2)-tA_str(1);    % calculate time difference between A1 and A2
    A2A1_cor=(A_crs(2)-A_str_bas)/(A_crs(1)-A_str_bas)*100;       % calculate the corrected A2/A1 index, from A1* and A2*
    A2A1=(A_str(2)-A_str_bas)/(A_str(1)-A_str_bas)*100;  %  calculate the orgional A2/A1 index, from A1* and A2*
else    % negative VGS
    d_tA_crs=tA_crs(2)-tA_crs(1);      % calculate time difference between A1 and A2
    A_crs(2)-A_crs_bas;
    A_crs(1)-A_crs_bas;
    A2A1_cor=(A_str(2)-A_crs_bas)/(A_str(1)-A_crs_bas)*100; % calculate the corrected A2/A1 index, from A1+ and A2+
    A2A1=(A_crs(2)-A_crs_bas)/(A_crs(1)-A_crs_bas)*100;  %   calculate the orgional A2/A1 index, from A1+ and A2+
end
% mark peaks on the plot
if length(A_crs)>=2
    % mark the A1, A2, A1* and A2*
    text(IDSCh1(TF_min,1),IDSCh1(TF_min,2),'\leftarrow A1, A2')
    text(IDSCh1(TF_max,1),IDSCh1(TF_max,2),'\leftarrow A1*, A2*')
else
    % mark the A for 1 pulse tests
    text(IDSCh1(TF_min,1),IDSCh1(TF_min,2),'\leftarrow A+')
    text(IDSCh1(TF_max,1),IDSCh1(TF_max,2),'\leftarrow A*')
end

% 2. weight change after last VGS change
% the lase A+ is the last change for VGS pulse
if PulDir
    LstVGSChgInd=find(IDSCh1(:,1)==tA_crs(PulNum));  % positive VGS, get 
    % the IDS array index for the last VGS change
    FstVGSChgInd=find(IDSCh1(:,1)==tA_str(1));  % positive VGS, get the IDS 
    % array index for the last VGS change


    A_PulDur(1,1)=tA_crs(1,1)-tA_str(1,1);    % get measured pules duration
    A_PulDur(2,1)=tA_crs(2,1)-tA_str(2,1);
    A_PulDet(1,1)=tA_str(2,1)-tA_crs(1,1);

else
    LstVGSChgInd=find(IDSCh1(:,1)==tA_str(PulNum));  % negative VGS, get 
    % the IDS array index for the last VGS change
    FstVGSChgInd=find(IDSCh1(:,1)==tA_crs(1));  % negative VGS, get the IDS 
    % array index for the last VGS change
    A_PulDur(1,1)=tA_str(1,1)-tA_crs(1,1);    % get measured pules duration
    A_PulDur(2,1)=tA_str(2,1)-tA_crs(2,1);
    A_PulDet(1,1)=tA_crs(2,1)-tA_str(1,1);
end

% initial IDS value
WghVGSChgInd=LstVGSChgInd+WghTimDly/SmpInt; % calculate the IDS array index 
% after weight change delay
%IDSWgh=IDSCh1(WghVGSChgInd,2);
IDSWgh=mean(IDSCh1(WghVGSChgInd-0.5*WghTimAvg/SmpInt:WghVGSChgInd+0.5*WghTimAvg/SmpInt,2));

%WghVGSIniInd=FstVGSChgInd-10*WghTimAvg/SmpInt;   % moved the averaging 
WghVGSIniInd=FstVGSChgInd-2*WghTimAvg/SmpInt;   % moved the averaging 
% window to 1 WghTimAvg before change point ot avoid including the change 
% points in the averaging value
IDSIni=mean(IDSCh1(WghVGSIniInd:FstVGSChgInd-WghTimAvg/SmpInt,2));

if PulDir   % positive VGS
    WghChgOut(1,1)=(A_str(2)-IDSIni)/(A_str(1)-IDSIni)*100;   % A2/A1*100%, Positive VGS pulses
    WghChgOut(2,1)=A_str(1);     % uA, IDS A1, positive pulses
    WghChgOut(3,1)=A_str(2);     % uA, IDS A2, positive pulses
else
    WghChgOut(1,1)=(A_crs(2)-IDSIni)/(A_crs(1)-IDSIni)*100;   % A2/A1*100%, negative VGS pulses
    WghChgOut(2,1)=A_crs(1);     % uA, IDS A1, positive pulses
    WghChgOut(3,1)=A_crs(2);     % uA, IDS A2, positive pulses
end
WghChgOut(4,1)=A_PulDur(1,1);   % s, VGS pulse duration, 1st 
WghChgOut(5,1)=A_PulDur(2,1);   % s, VGS pulse duration, 2nd 
WghChgOut(6,1)=A_PulDet(1,1);   % s, det time between pulses
WghChgOut(7,1)=IDSIni;          % uA, initial IDS
WghChgOut(8,1)=IDSWgh;          % uA, IDS at 30s after the last gate voltage change
WghChgOut(9,1)=(IDSWgh-IDSIni); % uA, det IDS 
WghChgOut(10,1)=(IDSWgh-IDSIni)/VDS % uS, det conductance 

% addtional IDS outputs
IDSCh1(:,4)=IDSCh1(:,2)-IDSIni;   % det IDS
% mark on plots
figure(1)
text(IDSCh1(WghVGSChgInd,1),IDSCh1(WghVGSChgInd,2),'\leftarrow Weight Change I_D_S')  %
text(IDSCh1(WghVGSIniInd+WghTimAvg/SmpInt,1),IDSCh1(WghVGSIniInd+WghTimAvg/SmpInt,2),'\leftarrow Initial I_D_S')
hold on

%% finishing plots
set(gca,'FontSize',16)
%legend('I_D_S (uA)')
figure(1)
title('Channel respond to gate voltage pulses')
xlabel('Time(s)')
ylabel('I_D_S (uA')
hold off

fprintf(IniNamStr1)
fprintf('done\n')
